# Machine Learning–Enhanced Nanopore ITS Analysis: Evaluating CPU–GPU Pipelines for High-Accuracy Fungal Taxonomic Resolution

**DOI:** 10.64898/2026.04.06.716835

**Authors:** Daniel Albuja, Peter Zambrano, Paolo Maldonado, J. Riczury Olmos-Tovar, Edwin Vera

## Abstract

Accurate fungal species identification is critical for microbial ecology, food safety, and plant pathology. However, morphological limitations and genomic complexity hinder this process. Molecular markers such as the ITS region, along with Oxford Nanopore long-read sequencing, offer a robust solution, albeit limited by error rates in homopolymeric regions and a high dependence on advanced computational resources (GPUs) to achieve high accuracy. This study benchmarks two bioinformatics workflows on a multiplexed dataset of complex fungal communities to address this technological gap: a CPU-based workflow optimized using a Bayesian machine learning engine and a GPU-accelerated workflow incorporating “super high accuracy” (SUP) models and refinement with neural networks. The results establish a scalable framework for evaluating the impact of computational architecture on final taxonomic resolution. It is demonstrated that GPU processing maximizes data retention and species-level accuracy by correcting systematic errors. Alternately, implementing automated hyperparameter optimization in CPU environments stabilizes sequence clustering and achieves high taxonomic concordance at the genus level. This conceptual advance validates the feasibility of performing ITS metabarcoding analysis in resource-constrained infrastructures, thus providing the scientific community with a reproducible protocol that balances the need for taxonomic precision with hardware availability.

## 1 Introduction

Fungal species identification is crucial for microbial ecology, plant pathology management, and agricultural biosecurity, as rapid diagnosis enables efficient countermeasures and minimizes economic losses (Loit et al., 2019; Yu et al., 2023). However, fungal identification is challenged by vast diversity and morphological limitations; diagnostic structures are often ephemeral, unstable, or morphologically identical across distinct taxa; hindering cryptic species (Borman et al., 2008; Nilsson et al., 2009).

To address these limitations, the scientific community has adopted molecular markers as standards for taxonomic classification (Schoch et al., 2012; Yu et al., 2023). Among these, the Internal Transcribed Spacer (ITS) region has been established as the universal barcode for fungi due to its high interspecific variability and technical advantages for PCR amplification (Nilsson et al., 2009). The ITS region is a segment of the nuclear ribosomal DNA (rDNA) cistron located between the Small Subunit (SSU or 18S) and Large Subunit (LSU or 28S) rRNA genes, comprising two highly variable non-coding spacer regions (ITS1 and ITS2) separated by the conserved 5.8S rRNA gene (Borman et al., 2008; Schoch et al., 2012). Its multi-copy genomic structure, with hundreds of tandem repeats per cell, facilitates PCR amplification even from samples with low initial DNA quantities (Nilsson et al., 2009).

The emergence of Oxford Nanopore Technologies (ONT) has enhanced ITS-based fungal identification by enabling full-length sequencing of the entire ribosomal operon (18S-ITS1-5.8S-ITS2-28S) in a single read, bypassing the fragmentation inherent to short-read platforms and improving phylogenetic resolution (Loit et al., 2019; Wang et al., 2021; Dierickx et al., 2024). However, ITS-based metabarcoding using nanopore sequencing presents inherent challenges. Length variability across taxa and the presence of homopolymeric regions, defined as consecutive repetitions of the same nucleotide, generate uniform electrical signals that complicate nucleotide inference, affecting basecalling precision and length estimation (Nilsson et al., 2009; Delahaye & Nicolas, 2021). Additionally, technical biases such as chimera formation, index switching, and preferential amplification of shorter fragments during PCR can distort community representation (Loit et al., 2019; Yu et al., 2023). These homopolymer-associated errors, including insertions, deletions (indels), and substitutions, result in higher error rates compared to short-read platforms, potentially compromising taxonomic assignment accuracy (Delahaye & Nicolas, 2021).

Advanced basecalling models, such as “super accuracy” (SUP), mitigate sequencing errors, substantially improving the resolution of complex genomic features (Purushothaman et al., 2025; Vereecke et al., 2025). However, achieving high precision through sophisticated models imposes a substantial computational burden. High-performance basecalling tools rely heavily on Graphics Processing Units (GPUs) with significant energy consumption to operate efficiently (Perešíni et al., 2021). Conversely, Central Processing Units (CPUs) often necessitate the use of lighter, less accurate models such as fast (FAST) basecalling, which can limit throughput or require prohibitive processing times (Ahsan et al., 2024; Delahaye & Nicolas, 2021). This technological disparity forces a trade-off: high-accuracy models on standard hardware are often prohibitively slow, requiring a compromise between quality and speed (Perešíni et al., 2021).

The quality of basecalled sequences is a critical determinant for downstream processing and taxonomic assignment. Sequence clustering has emerged as an essential post-processing step to further improve data quality by aggregating DNA sequences into Operational Taxonomic Units (OTUs) based on similarity thresholds, typically 98% for species-level differentiation (Loit et al., 2019; Dierickx et al., 2024). This process serves dual purposes: correcting residual sequencing errors through consensus sequence generation and reducing computational complexity by collapsing highly similar reads. However, the effectiveness of clustering is contingent upon initial read quality, as high error rates can generate spurious clusters and inflate diversity estimates, particularly for complex markers like fungal ITS (Loit et al., 2019; Dierickx et al., 2024). Frameworks such as VSEARCH are frequently employed for amplicon clustering in nanopore workflows (Dierickx et al., 2024). Nevertheless, advanced consensus generation tools often require manual hyperparameter tuning, rendering the process subjective and laborious (Dierickx et al., 2024).

Despite advancements in sequencing technologies, notable gaps remain in the optimization of bioinformatic workflows for fungal ITS metabarcoding. To our knowledge, machine learning-driven hyperparameter optimization has not been systematically evaluated for nanopore ITS sequence clustering. Although computational performance comparisons between CPU and GPU architectures exist for basecalling and methylation detection (Perešíni et al., 2021; Ahsan et al., 2024), direct assessments of their impact on final taxonomic accuracy in ITS analysis remain limited. Most available nanopore ITS workflows focus predominantly on benchmarks against Illumina or the performance of a single workflow using individual isolates or simulated communities of limited complexity (Loit et al., 2019; Dierickx et al., 2024; Purushothaman et al., 2025).

To overcome these gaps this study benchmarks two complete bioinformatic workflows for processing nanopore ITS amplicons: a CPU-based pipeline incorporating machine learning-driven hyperparameter optimization via Optuna, and a GPU-accelerated pipeline integrating SUP basecalling models with neural network-based consensus polishing. Using a multiplexed dataset of 28 barcoded samples representing complex fungal communities with known expected taxa, we established a scalable framework for evaluating the impact of computational architecture on taxonomic resolution. The integration of Bayesian optimization automated parameter selection, enhancing reproducibility and reducing subjective bias inherent in manual tuning. By systematically comparing both workflows, this study provides practical guidance for selecting analysis strategies based on accuracy requirements and available computational resources, thereby facilitating more accessible and robust pipelines for fungal identification in plant pathology, food safety, and environmental surveillance applications.

## 2 Materials and Methods

### 2.1 Sample Collection and Biological Context

Fungal isolates were obtained from banana, pitahaya, and pineapple peels that had been kept under natural ambient conditions in the postharvest laboratory in Quito, where the mean annual temperature is about 14 °C and the relative humidity is approximately 83% (INAMHI et al., 2018; Weather Atlas, 2022). The samples were supplied by the Postharvest Laboratory of the Department of Food Science and Biotechnology (DECAB) at the Escuela Politécnica Nacional (EPN). Prior to this study, the isolates had been purified as monosporic cultures and preliminarily identified at DECAB through Sanger sequencing of the Internal Transcribed Spacer (ITS) region.

#### 2.1.1 DNA Extraction and ITS Amplification

DNA was extracted from seven-day-old fungal cultures using a protocol adapted from Cenis (1992). Prior to extraction, monosporic cultures were grown on PDA for 7 days at 28 °C. Lysis was performed by vortexing 400 µL of lysis buffer, 4 mm glass beads, and a ∼2 mm mycelial plug in a 2 mL microcentrifuge tube for 20 min. Then, 200 µL of 3 M sodium acetate (pH 5.2) was added, mixed briefly, and the tube was incubated at −20 °C for 20 min. Samples were centrifuged at 14,000 rpm for 20 min at room temperature, and the supernatant (≈500 µL) was transferred to a new nuclease-free tube. An equal volume of isopropanol was added to precipitate DNA for 2 hours at ambient conditions, followed by centrifugation at 14,000 rpm for 10 min. The pellet was washed with 70% ethanol twice, air-dried overnight, and resuspended in 30 µL of nuclease-free water. DNA concentration and purity were determined using a NanoDrop One spectrophotometer. Extraction was performed in two biological replicates.

Amplification of the ITS region was conducted using the universal primers ITS1 (5’-TCCGTAGGTGAACCTGCGG-3’) and ITS4 (5’-TCCTCCGCTTATTGATATGC-3’) obtained from Integrated DNA Technologies (IDT). PCR reactions were run under the following thermal program: an initial denaturation at 95 °C for 5 min; 34 cycles of 95 °C for 30 s, annealing at 52 °C for 1 min, and extension at 72 °C for 2 min; and a final elongation step at 72 °C for 10 min. Amplicon quality was verified by electrophoresis on a 1.5% agarose gel. PCR products were purified using the Mag-Bind Total Pure NGS Kit (OMEGA BIO-TEK, 2018) in accordance with manufacturer’s protocol and quantified using a Qubit 4 fluorometer.

#### 2.1.2 Library preparation and Sequencing conditions

DNA sequencing was carried out using the Rapid Barcode g-DNA Sequencing Kit (Cat. No. RBK11096 10.0017; SQK-RBK96) from Oxford Nanopore Technologies (2023), following the manufacturer’s protocol. Sequencing runs were performed on a MinION Mk1C instrument equipped with R9.4.1 flow cells, with a 24-hour runtime, and each sample was processed duplicate.

### 2.2 Post-processing

Two complete ITS post-processing pipelines were implemented to evaluate the influence of computational architecture on sequence accuracy, clustering behavior, and taxonomic resolution. Both workflows began with raw FAST5 nanopore reads and terminated in chimera-filtered consensus taxonomic assignment tables. The CPU implementation was based on a traditional sequential processing framework (Figure M1), whereas the GPU pipeline integrated an amplicon grouping approach and a deep learning-driven consensus refinement step (Figure M1). The two pipelines were maintained parallel in structure to enable controlled comparison using identical input samples comprising 28 barcoded libraries.

**Figure M1.**
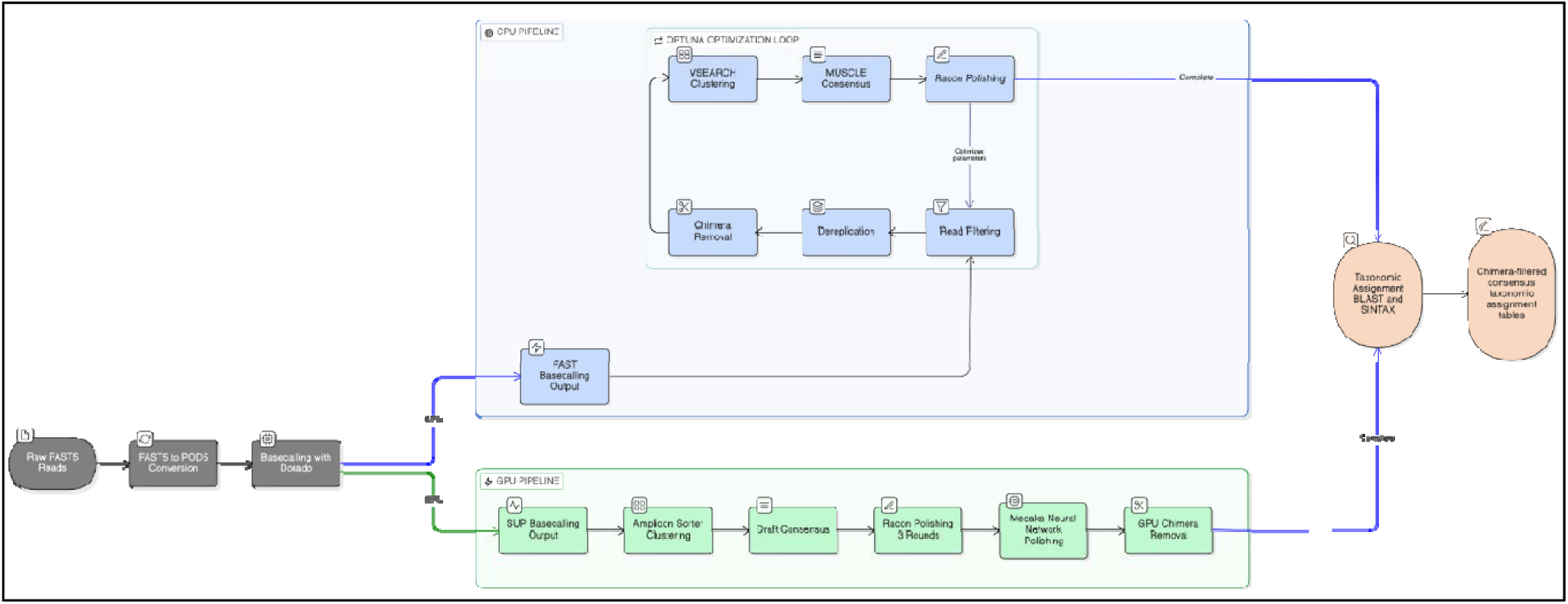
Overview of the CPU and GPU ITS processing workflows

#### 2.2.1 Dorado basecalling

Dorado is an Oxford Nanopore Technologies’ open-source, basecalling software that converts raw electrical current measurements from nanopore pores into nucleotide sequences using deep neural network architectures. It supersedes Guppy as ONT’s primary recommended basecaller and provides configurable accuracy tiers (FAST, HAC (high accuracy), and SUP(super accuracy)), allowing users to balance computational throughput with sequence accuracy according to available hardware resources (Purushothaman et al., 2025).

Raw nanopore electrical signals were processed independently for each barcode using a barcode-by-barcode processing strategy. Automated scripts sequentially executed each post-processing step on demultiplexed reads for individual barcodes.

Because Dorado (v0.9.6) requires POD5 as its preferred input format, each barcode directory was processed through a two-stage signal conversion workflow. Individual FAST5 files were converted to POD5 format using the official ONT pod5 library (v0.3.34) executed via Python. All POD5 files generated per barcode were subsequently merged into a single multi-read POD5 file using pod5_merge.py. Failed FAST5 to POD5 conversions were automatically logged, and only valid merged POD5 files were preserved. Intermediate POD5 fragments were removed to reduce storage overhead.

Basecalling was conducted using Dorado v0.9.6, the higher version compatible with the R9.4.1 flow cell available at the time of analysis. All basecalling runs were performed in simplex mode, as duplex reads were not available in the raw sequencing output.

Each barcode was basecalled independently, exporting final reads in FASTQ format. A strict minimum Q-score threshold of 10 (--min-qscore 10) was applied uniformly to both FAST and SUP basecalling models. To quantify performance differences due to model complexity, two Dorado configurations were evaluated:

##### CPU Workflow

␿ *Model:* dna_r9.4.1_e8_fast@v3.4
␿ *Hardware:* Google Colaboratory (CPU-only environment)

The FAST@v3.4 model was selected for the CPU workflow because it represents the highest-performance Dorado architecture capable of operating within realistic CPU-only computational constraints. According to Peresini et al. (2021), more accurate models like HAC or SUP do not reach practical runtime on CPU-only systems and require GPU acceleration.

##### GPU Workflow

␿ *Model:* dna_r9.4.1_e8_sup@v3.3
␿ *Hardware:* CEDIA High Performance Computing (HPC) with GPU acceleration

For the GPU workflow, the SUP@v3.3 model was selected because it maximizes read-level Q-scores, reduces indel frequency in homopolymeric regions, and improves downstream clustering stability, all of which are critical requirements for resolving ITS-level fungal diversity (Dierickx et al., 2024).

#### 2.2.2 Adapter trimming (Porechop)

Adapter trimming was performed using Porechop_ABI, (v0.5.0), which identifies and removes adapter sequences from raw FASTQ reads but retains barcode information. This tool was selected due to its proven efficiency with long-read data and its ability to discard internal adapter sequences that may interfere with downstream clustering, as described in Bonenfant et al. (2023, p. 1).

Trimming was performed in a dedicated Conda environment on Google Colaboratory (Ubuntu Linux x86_64), using two threads for parallel processing to optimize computational efficiency. For each barcode directory, adapter trimming was applied to the corresponding single FASTQ file generated during basecalling. The –discard_middle option was explicitly activated to discard reads containing internal adapters sequences, while all other parameters were left at their default settings. No explicit minimum read length threshold was set; therefore, reads reduced to 1 bp or shorter during trimming were discarded according to Porechop’s default behavior. Barcode removal was not enforced at this stage, as barcodes had already been removed during basecalling-based demultiplexing; however, Porechop automatically screened for potential residual barcode sequence matches.

For each barcode, the pipeline logged the read count prior to trimming and generated a dedicated log file containing all execution outputs, error messages and process return codes. This ensured full traceability and reproducibility of the trimming step.

#### 2.2.3 Pipeline 1: CPU-based Workflow

##### 2.2.3.1 Clustering with VSEARCH

ITS read clustering was done using VSEARCH (v2.23.0) within a Python-based automated optimization pipeline. VSEARCH was selected because it is a versatile, open-source metagenomics tool that supports amplicon dereplication, and clustering, is efficient for long-read Oxford Nanopore data, and integrates UCHIME-based de novo chimera detection (Rognes et al., 2016).

For each barcode-specific FASTQ file resulting from trimming, an automated script executed a multi-step workflow designed to maximize cluster stability and consensus sequence quality. The pipeline was executed on Google Colaboratory (Ubuntu Linux x86_64) and parallelized across optimization trials using Optuna. Every optimization trial entailed performing the following steps:

1. **Read filtering.** Raw reads were filtered based on a minimum read length and a minimum mean Phred quality score, both treated as tunable hyperparameters.
2. **Dereplication.** Filtered sequences were dereplicated at 100% identity using VSEARCH (--derep_fulllength –sizeout), collapsing identical full-length reads while retaining their abundance information to reduce redundancy and improve computational efficiency in downstream analyses.
3. **Chimera detection.** De novo chimera removal was performed with UCHIME-denovo (--uchime_denovo) to artifactual chimeric sequences prior to OTU (Operational Taxonomic Unit) formation.
4. **Clustering.** Non-chimeric sequences were clustered using VSEARCH’s size-based clustering algorithm (--cluster-size), which prioritizes high-abundance sequences, with bidirectional search enabled (--strand both). The clustering identity threshold (--id) was optimized by Optuna. VSEARCH produced both centroid sequences and a .uc cluster mapping file.
5. **Consensus generation.** For each cluster meeting a minimum cluster size, a multiple sequence alignment was computed with MUSCLE (v3.8.31). Consensus sequences were generated using a custom Python module applying a majority-vote rule, with the identity threshold treated as an additional hyperparameter optimized by Optuna.
6. **Polishing.** Consensus sequences were polished using a single round of Racon, balancing error correction with the risk of over-polishing.

For each trial, all intermediate files were saved, including filtered reads, dereplicated sequences, centroids, .uc mapping files, consensus sequences and polished outputs. After optimization, the polished consensus sequences from the best-performing trial were selected for downstream taxonomic analysis.

All VSEARCH, MUSCLE, Minimap2, and Racon commands, as well as Optuna search spaces and trials metrics, were logged automatically to guarantee full reproducibility.

##### 2.2.3.2 Optuna hyperparameter optimization

To improve the performance and robustness of the clustering workflow, an automated hyperparameter optimization strategy was implemented using Optuna (v3.6.1), an open-source framework for efficient hyperparameter tuning (Akiba et al., 2019). This optimization was performed independently for each barcode and sought to find the combination of parameters that maximize cluster coherence, minimize spurious OTUs, and improve the quality and reliability of the final consensus sequences.

Five hyperparameters were optimized to account for the methodological heterogeneity reported for fungal ITS metabarcoding studies using Nanopore sequencing and VSEARCH-based clustering, where the parameter choices depend substantially on the amplicon length, sequencing error profile, and taxonomic resolution desired:

1. Minimum read length (range: 400-1000 bp)
2. Minimum mean Phred quality (range: 10-20)
3. VSEARCH clustering identity threshold (--id, range: 0.85-0.99)
4. Minimum cluster size required to generate a consensus (range: 2-5 reads)
5. Consensus identity threshold for base calling within MUSCLE alignments (range: 0.4-0.8)

Each Optuna trial executed the full clustering pipeline described in Section 2.2.3.1 At the end of each Optuna trial, a single composite objective score was computed and minimized to guide hyperparameter selection. The objective function was explicitly defined as the sum of multiple penalty terms capturing sequence quality, clustering stability, and taxonomic reliability:

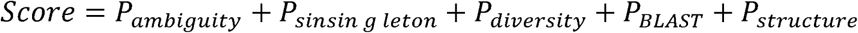

Lower scores indicate higher-quality clustering and consensus outcomes.

The ambiguity penalty corresponds to the average percentage of ambiguous bases (N) across polished consensus sequences, reflecting uncertainty in base calling. The singleton penalty is defined as the proportion of single-read clusters relative to the total number of clusters, scaled to penalize spurious or weakly supported OTUs. The diversity penalty is derived from the normalized Shannon entropy of the cluster size distribution, penalizing excessive fragmentation relative to the maximum achievable entropy for a given number of clusters.

Taxonomic reliability was incorporated through a BLAST-based penalty, computed as a smooth logarithmic function of deviations from perfect sequence identity and alignment coverage, thereby avoiding extreme score inflation while still discouraging poorly supported consensus sequences. Finally, a structural penalty was applied when fewer than three clusters contained more than one read, discouraging under-clustering scenarios dominated by insufficiently supported consensus sequences.

To determine an appropriate number of trials, a convergence analysis was performed using representative barcodes from each main fungal group present in the dataset. Within each group, the representative barcode was defined as the sample whose read count was closest to the group median. Pilot optimization runs were executed for these representative barcodes and their full Optuna histories exported. For each run, the cumulative best score was tracked across trials while computing the marginal improvement over a sliding window of five consecutive trials. A window size of five was chosen to smooth short-term stochastic fluctuations inherent in Bayesian optimization while preserving sensitivity to systematic performance gains across iterations. An improvement threshold of ε = 1% was defined as the minimum change considered meaningful.

The median marginal improvement curve across the three representative barcodes stabilized below ε after about 12 trials, which means additional trials produced negligible improvements in the objective function. Finally, 12 trials per barcode were selected for the full CPU workflow as a balance between computational cost and convergence stability. All optimization histories and best-performing parameter sets were exported. The polished consensus associated with the best trial was saved and used for downstream taxonomic assignments.

#### 2.2.4 Pipeline 2: GPU-based Workflow

##### 2.2.4.1 Amplicon Sorter grouping strategy

Amplicon-level read grouping in the GPU workflow was performed using Amplicon Sorter (commit corresponding to the December 2025 release available at the time of analysis), a tool specifically designed to cluster long Oxford Nanopore reads into amplicon-consistent groups prior to consensus generation (Vierstraete & Braeckman, 2022). Amplicon Sorter combines GPU-accelerated similarity scoring with iterative clustering strategies optimized for high-error long-read data, improving the recovery of low-abundance ITS variants relative to conventional CPU-based clustering approaches.

For each barcode, trimmed FASTQ files generated by Porechop were processed independently in single-sample mode, as recommended for barcode-resolved amplicon datasets. A sequence similarity cutoff of 90% was applied during clustering, following the developer’s recommendations for fungal ITS amplicons sequenced with ONT R10 chemistry. All other parameters were kept at their default values.

Depending on data quality and read support, two alternative outcomes were possible. In the typical case, Amplicon Sorter successfully completed both the grouping and internal polishing steps, resulting in a draft consensus sequence that was retained for downstream polishing and comparison steps as the initial amplicon-level reference. In a small number of barcodes, Amplicon Sorter failed due to insufficient read overlap, low sequencing depth, or excessive read fragmentation. In these cases, a fallback rescue pathway was invoked. In this procedure, reads were aligned using Minimap2 (v2.30, ONT presets), and a single-round consensus was generated using Racon (v1.5) to recover a provisional draft sequence. If successful, this rescue consensus was used as the initial draft in subsequent analyses. Barcodes for which neither the primary nor the rescue procedure yielded a valid draft were retained in the dataset but were explicitly flagged as lacking a usable consensus.

The dual-strategy design was employed to ensure consistent treatment across all barcodes while minimizing data loss due to clustering instabilities. The rescue pathway was invoked only in rare cases and served to maintain comparability across barcodes during downstream polishing and quality assessment.

##### 2.2.4.2 Medaka and Racon polishing

In the GPU-based workflow, consensus refinement was performed using Racon correction followed by Medaka neural-network polishing.

Racon polishing was initiated from the draft consensus sequence. For each barcode, raw Nanopore reads were used as polishing input. Three iterative Racon rounds were executed to progressively reduce indel errors in ITS amplicons. Each iteration consisted of read-to-consensus alignment using Minimap2, followed by error correction with Racon (v1.5.0). For each iteration, alignments were generated in PAF format and passed directly to Racon, with both mapping and polishing executed using two threads.

Following Racon correction, Medaka (version 1.11.3) was used to carry out refinement of the consensus using the r941_min_sup_g507 model, chosen for compatibility with R9.4.1 chemistry. This tool uses deep neural networks models to compute high-accuracy consensus sequences from aligned reads, improving sequence quality for accurate taxonomic classification in fungal ITS amplicons (Lee et al., 2021). Medaka polishing was run within a dedicated Conda environment with GPU acceleration enabled. It produced corrected consensus sequences and variant files for each barcode.

The combined Racon-Medaka polishing strategy reduced homopolymer-associated errors typical of Oxford Nanopore reads, producing final consensus sequences suitable for downstream taxonomic classification.

##### 2.2.4.3 Chimera detection

To identify and remove any PCR-derived artefacts that may artificially inflate diversity estimates in amplicon-based fungal metabarcoding, chimera detection was performed on polished consensus sequences (Hakimzadeh et al., 2025). This step was performed using VSEARCH (v2.23.0) and was applied only to barcodes for which a final Medaka-polished consensus sequence was successfully generated.

A dereplication-first strategy was applied to improve both computational efficiency and chimera detection accuracy. In the first stage, full-length dereplication collapsed identical consensus sequences while retaining abundance information. This reduces redundancy among consensus sequences and improves the robustness of downstream chimera detection by focusing on unique variants.

The dereplicated sequences were then subjected to de novo chimera detection using the UCHIME algorithm. A de novo approach was selected to avoid database-dependent biases and because extensive error correction had already been applied earlier in the GPU workflow. UCHIME de novo classified sequences into putative chimeric and non-chimeric sets based on internal self-consistency modeling.

Resulting files were stored in dedicated subdirectories within each barcode folder to ensure clear separation from upstream processing stages and full reproducibility. Processing logs were retained for all analyzed barcodes. This final quality control step removed artefactual consensus sequences prior to taxonomic assignment and harmonized chimera handling between the CPU- and GPU-based pipelines.

#### 2.2.5 Taxonomic assignment

A hybrid pipeline that combines BLASTn (version 2.17.0) with SINTAX implemented in VSEARCH (version 2.23.0) (Altschul et al., 1990; Rognes et al., 2016) performed taxonomic assignment of the polished consensus sequences. All consensus sequences generated during the VSEARCH step were aggregated into a single multi-FASTA file. Each sequence ID was modified to encode its barcode of origin, ensuring traceability during downstream taxonomy operations.

BLAST searches were run against a custom UNITE database (unite2024ITS) created from the 2024 UNITE release, selected due to its comprehensive coverage of fungal ITS sequences (Abarenkov et al., 2024). For correct genus and species parsing, an accession-to-taxonomy map was compiled via scanning all headers in UNITE_public_19.02.2025.fasta, corresponding to the UNITE 2024 release and extracting standard UNITE tokens.

BLAST searches were performed using tabular output format with extended fields, such as query coverage per subject, query coverage per HSP, and subject title. The number of reported hits was limited to 10 per query. An e-value threshold of 1e−5 was applied. An early permissive filter was applied to discard trivial or spurious matches: only hits with qcovhsp > 1% were retained at this stage.

SINTAX taxonomic classification was performed using the UNITE UTAX reference. The pipeline parsed genus and species level annotations along with their SINTAX confidence scores. All results were exported and retained without pre-filtering.

BLAST and SINTAX outputs were integrated using a hierarchical set of deterministic decision rules.

␿ **Rule A: High-confidence species (SINTAX + BLAST agreement)** - A SINTAX species confidence threshold of 0.9 was adopted. BLAST-based species assignment was constrained by using a minimum percent identity of 97% and coverage threshold of 90%. Species-level assignments were accepted only when SINTAX and BLAST converged to the same species as the top BLAST hit.

␿ **Rule B:** Extreme BLAST dominance - A conservative threshold for BLAST identity ≥99.0% was set, with a high alignment coverage requirement (≥95%). A minimum percent identity difference (≥1.0%) between the top and second BLAST hits was required.

␿ **Rule C:** Genus-level consensus - A BLAST percent identity threshold of ≥94.0% was used, with a minimum BLAST query coverage of approximately 80%. A stricter SINTAX genus confidence threshold of ≥0.90 was applied. Only cases in which the genus inferred by SINTAX matched the genus derived from BLAST were retained under this rule.

If none of the rules A-C were satisfied, additional fallbacks were applied, including Least Common Ancestor (LCA) inference across all BLAST hits, a weak-hit genus-level fallback when bitscore values exceeded 100 and genus names were consistent, and a default assignment of “Fungi sp.” When no reliable taxonomy could be inferred.

Queries that did not meet any threshold or fallback rule were flagged as unassigned.

A composite penalty score was computed for each assigned OTU to quantify assignment uncertainty:

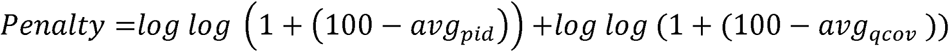

where avgpid is the average percentage identity from BLAST, and avgqcov is the average query coverage.

The pipeline produced three structured tabular outputs. The main output includes one row per OTU with the assigned taxonomic rank and name, the applied decision rule, associated BLAST and SINTAX metrics, and the composite penalty score. Finally, another .tsv summarizes OTUs for which no reliable taxonomic assignment could be made, reporting insufficient or conflicting taxonomic signals together.

#### 2.2.6 Data and Code Availability Statement

The code used in this study is publicly available at GitHub: https://github.com/sequencingnanopore-create/ITS-nanopore-cpu-gpu-benchmark. A permanent archived version of the repository is available on Zenodo: https://doi.org/10.5281/zenodo.19325905. All scripts, notebooks, and documentation required to reproduce the analyses are included. The computational pipeline used in this study is an adapted and optimized version of an original workflow developed in Google Colab. The code was modified to improve reproducibility, portability, and accessibility, enabling execution across different local computing environments without reliance on cloud-based platforms.

Raw sequencing data (FASTQ files generated using the CPU-based basecalling pipeline) have been deposited in the NCBI Sequence Read Archive (SRA) under BioProject accession PRJNA1444816. These data correspond to the primary dataset used in the analyses presented in this study.

## 3 Results

### 3.1 Dataset Overview and Initial Read Characteristics

Table R1 summarizes the initial data processing metrics across 28 multiplexed fungal barcodes, specifically focusing on the results up to the trimming stage. It compares the read retention between the CPU-based workflow and the GPU-based workflow. These preliminary statistics establish the baseline data quantity available for subsequent clustering and taxonomic analysis.

**Table R1.**
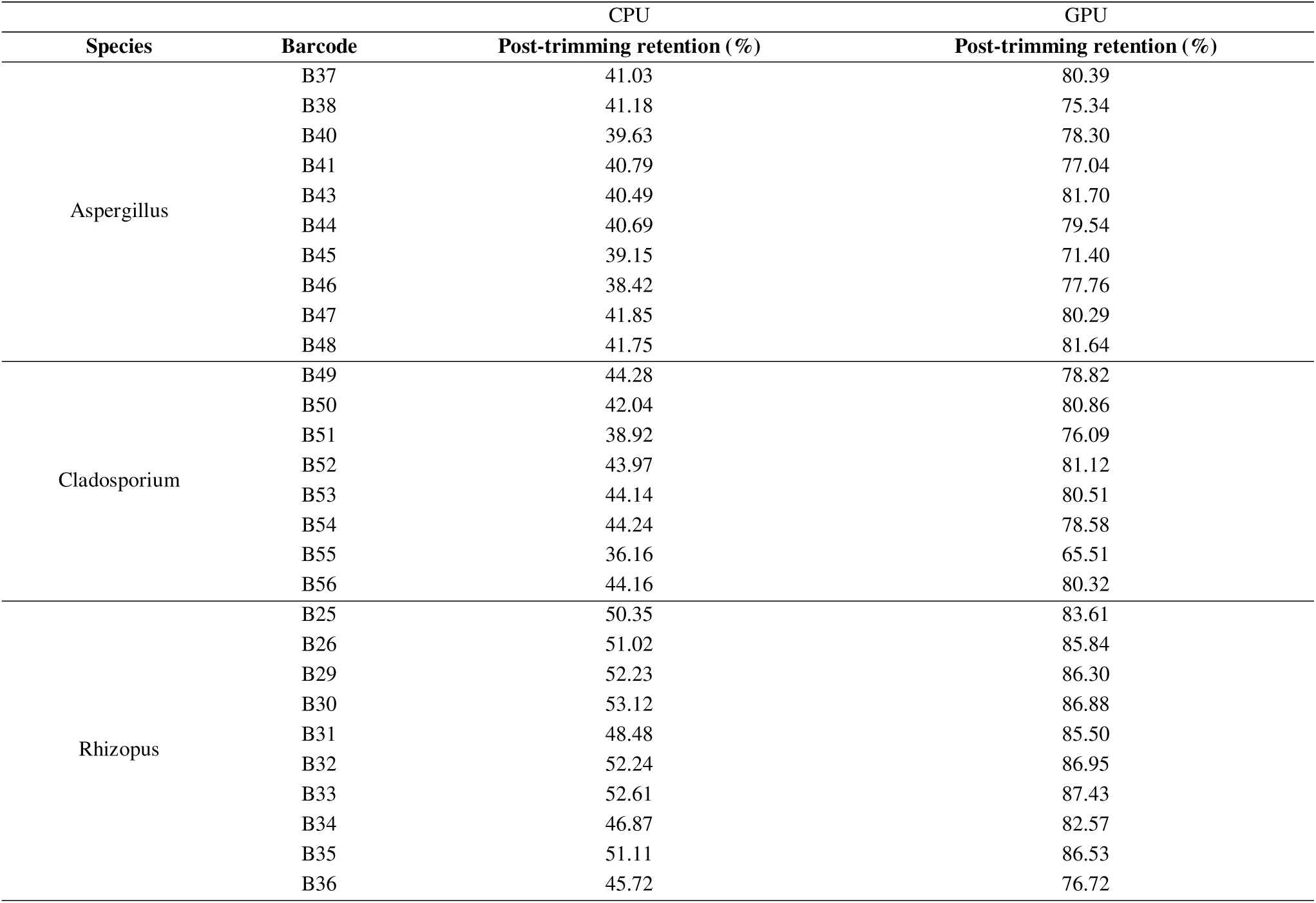
Summary of read counts and length distributions across expected-taxon groups.

Figure R1 shows the read density of data retrieval. It can be observed that the distributions corresponding to GPU processing exhibit a higher median and a wider probability density toward high read count values compared to CPU processing for the three fungal genera studied (Aspergillus, Cladosporium, and Rhizopus).

**Figure R1.**
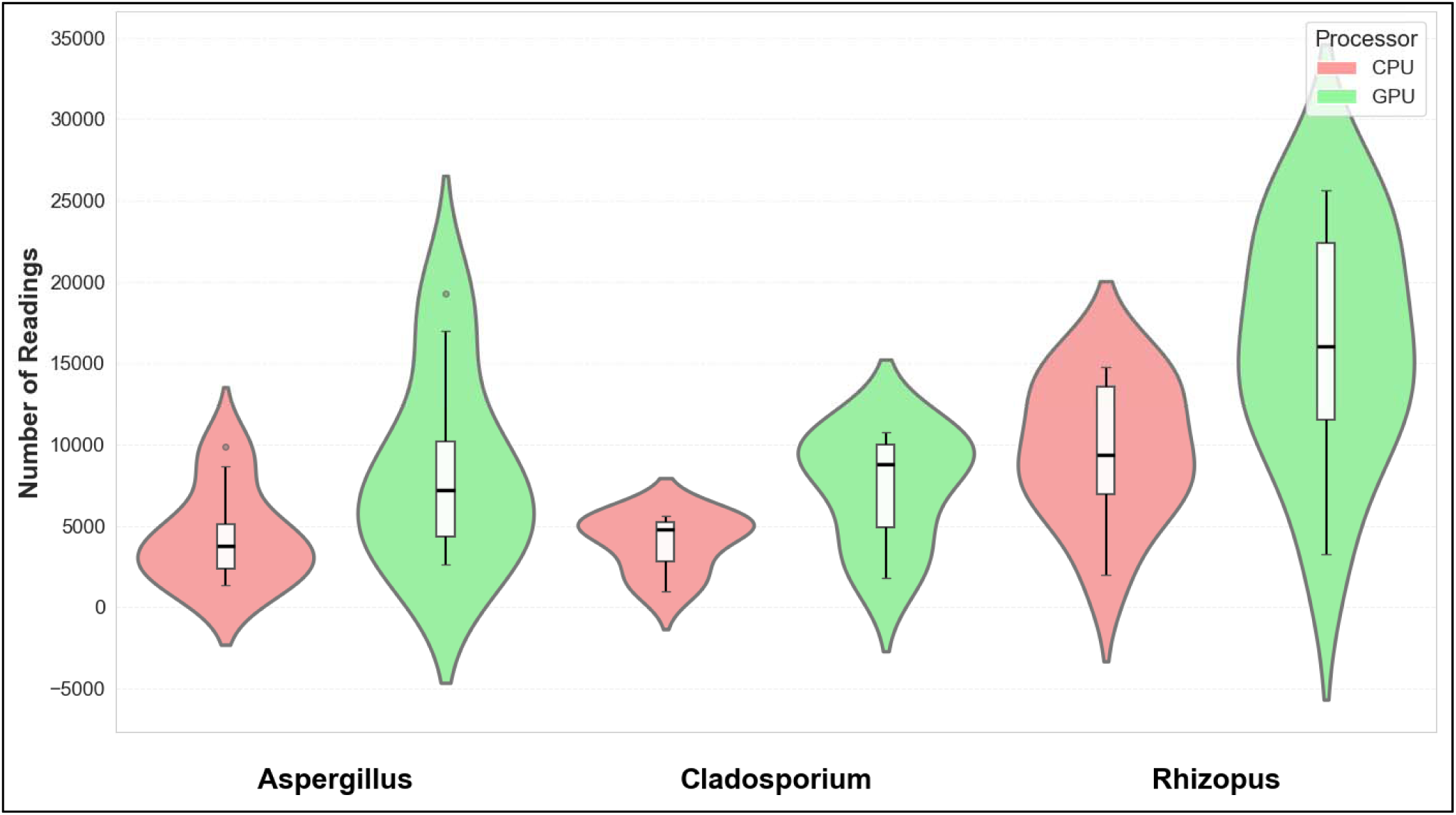
Read count distributions per group (CPU vs GPU porechop)

### 3.2 Basecalling Performance (FAST vs SUP)

**Figure R2 (A).**
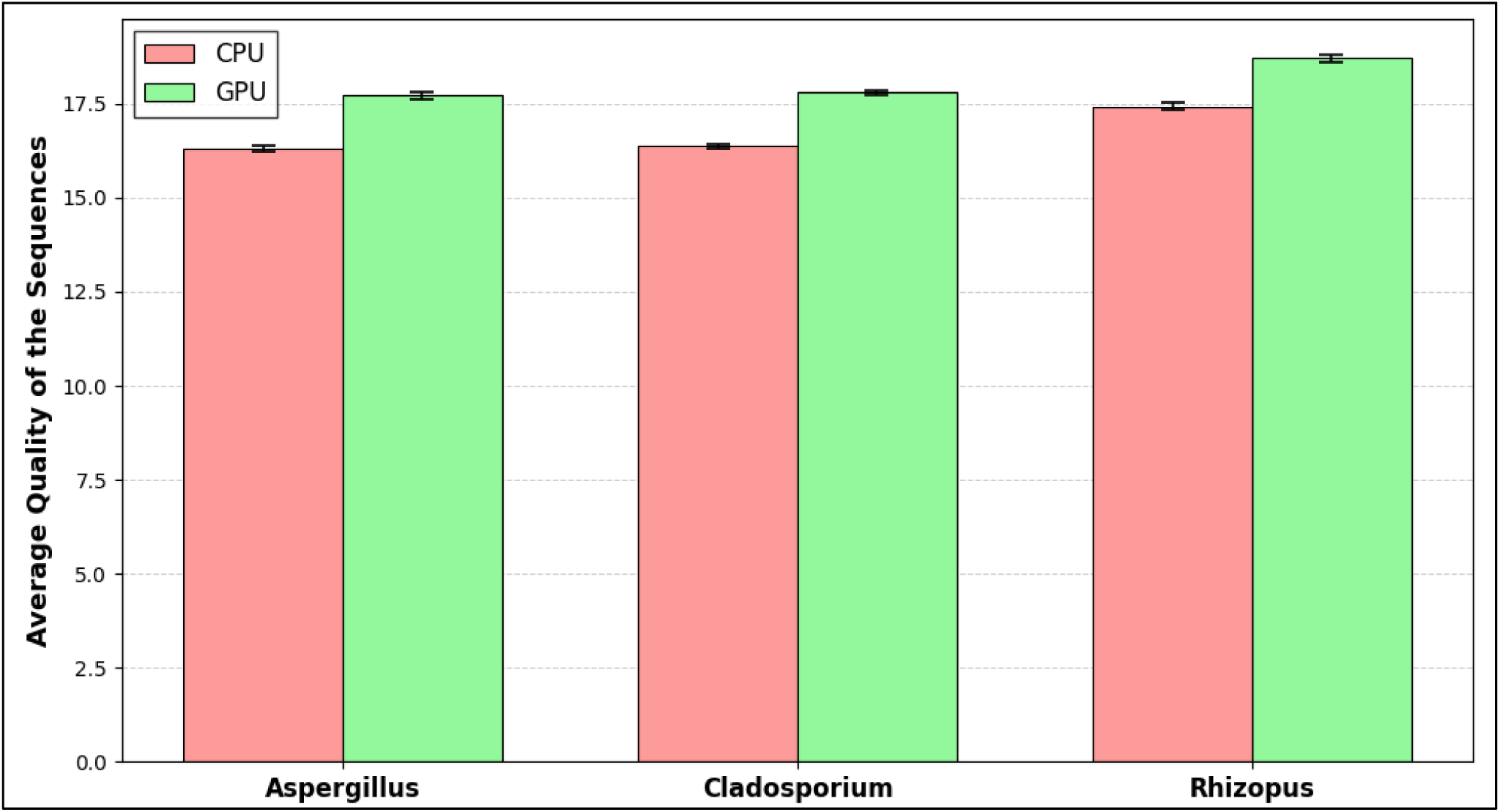
Basecalled read quality metrics, with Q-scores per group/model

Analysis of the quality metrics of the generated sequences shows an improvement in mean quality scores (Q-scores) when using GPU-based workflows compared to CPU processing. As shown in Figure R2(A), the Phred scores obtained using the high precision basecalling (SUP) model on GPUs outperform those of the fast (FAST) model on CPUs for all three fungal genera evaluated.

**Figure R2 (B).**
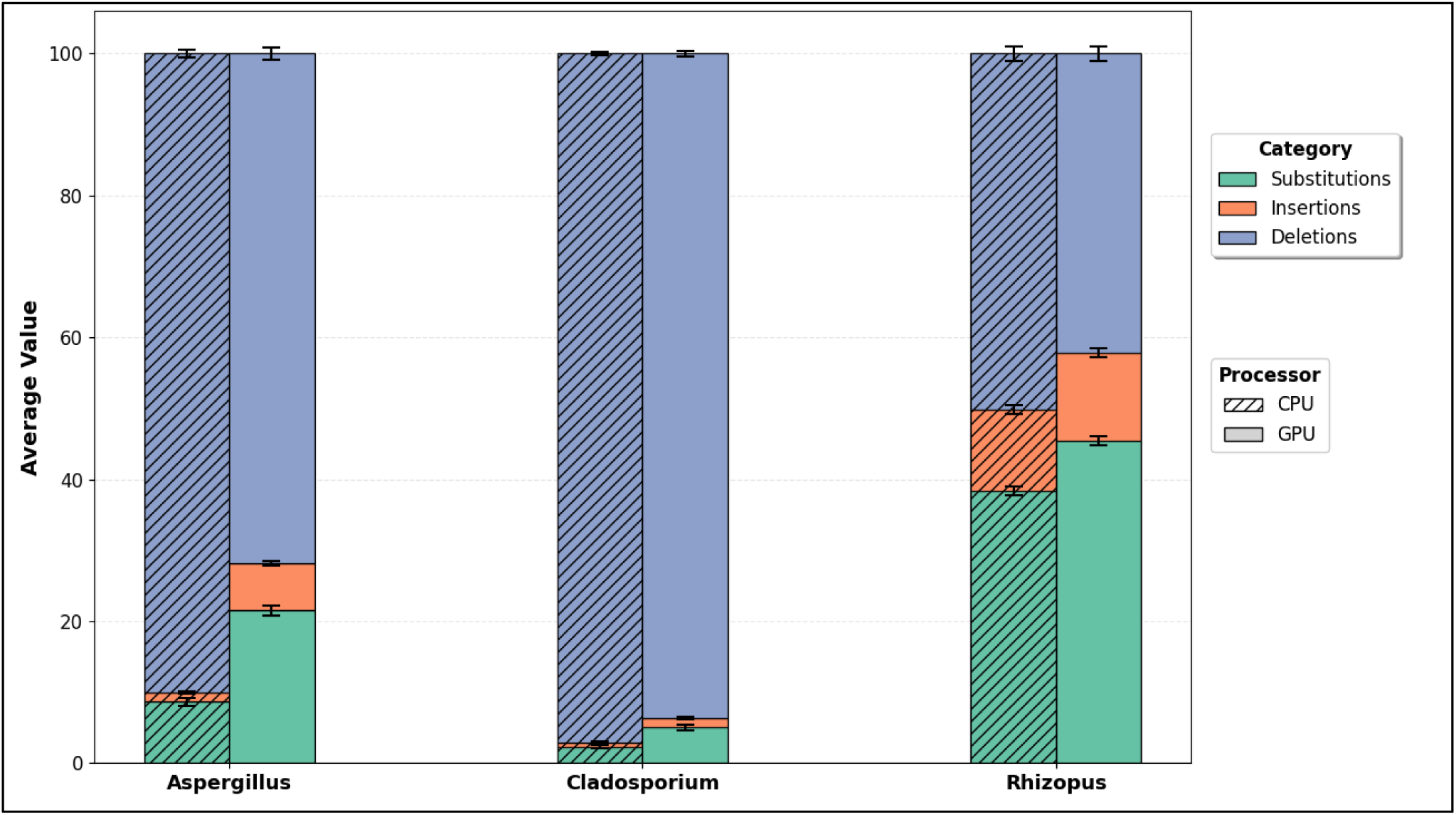
Basecalled read quality metrics, Stacked error bars

In the CPU-executed workflow, Figure R2 (B) shows deletions predominate over insertions and substitutions, constituting most of the total error load, particularly in the genera Aspergillus and Cladosporium. This pattern demonstrates the limitations of fast basecalling (FAST) models, which lack the computational complexity necessary to adequately resolve long homopolymeric regions. Consequently, in these areas, the nanopore’s electrical signal remains static, and simplified models tend to underestimate the event’s duration, resulting in the systematic omission of nucleotides in the generated sequence (Delahaye & Nicolas, 2021). Thus, the elevated error burden observed in the CPU-based workflow, amounting to 1,183,250 total errors across 8,400 reads, indicates that speed-oriented basecalling compromises amplicon structural integrity, introducing length biases that can negatively affect alignment accuracy and downstream taxonomic classification.

Conversely, implementation of the pipeline on GPUs using the high-precision (SUP) model substantially alters the error profile. Although the GPU-based workflow exhibits a slightly higher overall number of errors (1,317,654 total errors across the same number of reads), the dominance of systematic deletion artifacts is reduced, resulting in a more balanced composition in which residual errors are predominantly stochastic in nature. This shift does not imply a deterioration in basecalling accuracy but rather reflects the enhanced capacity of SUP neural network architectures to correct systematic deletion events characteristic of FAST models, leaving less structured errors as the primary residual class (Wick et al., 2019). Ultimately, the higher prevalence of errors in the CPU-based workflow underscores the critical necessity of deploying more complex, GPU-accelerated processing pipelines to mitigate these systematic artifacts and ensure data fidelity for accurate fungal identification (Purushothaman et al., 2025; Wang et al., 2021).

### 3.3 CPU Pipeline Results

#### 3.3.1 Optuna Optimization Behavior

The behavior of the Optuna-based optimization process was evaluated across all barcoded samples, with three representative barcodes (B34, B41 and B56) selected to illustrate the structure of the optimization landscape, as shown in Figure R3.

**Figure R3.**
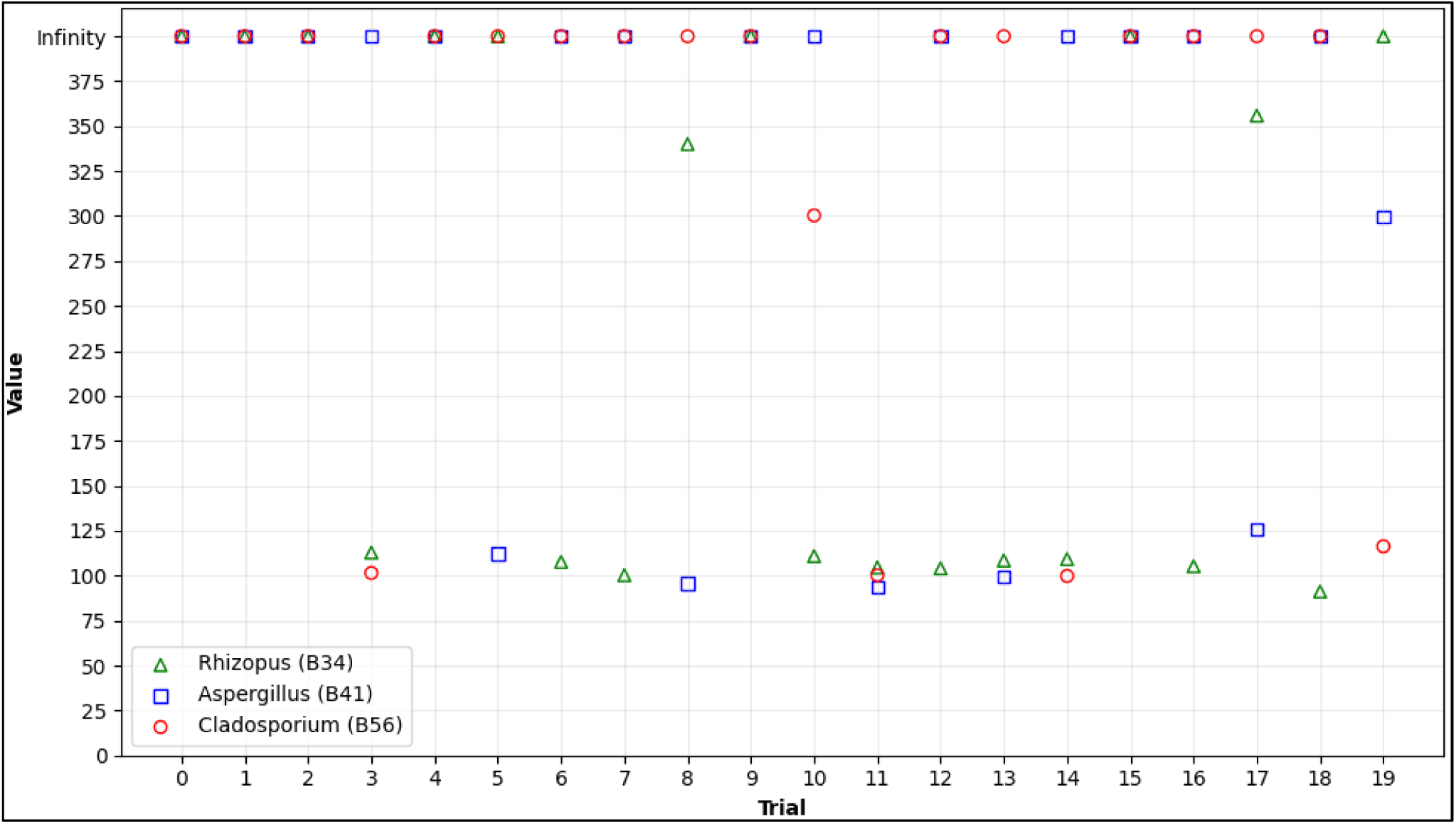
Optuna search landscape

Across all three barcodes, a substantial fraction of trials resulted in infeasible outcomes, defined as parameter combinations that produced no valid consensus sequences after polishing and downstream filtering. These trials were deliberately assigned an infinite objective value within the optimization framework and are explicitly represented in the optimization landscape depicted in Figure R3. The frequency of infeasible trials varied among barcodes, with B41 exhibiting the highest proportion of infinite values, B56 an intermediate proportion, and B34 the largest number of trials yielding finite objective values.

For trials producing finite outcomes, objective values displayed distinct barcode-specific distributions over the course of the optimization. Barcode 56 achieved the lowest objective value overall, reaching a minimum of approximately 88 at trial 7. Barcode B34 reached its minimum objective value of approximately 97, also at trial 7, and exhibited the widest range of finite objective values across trials, including isolated high-value outcomes exceeding 300. In contrast, barcode B41 attained its lowest objective value, approximately 101, earlier in the optimization process at trial 3 and showed a comparatively narrow distribution of objective values across feasible trials.

Across all three barcodes, low objective values were not confined to a single stage of optimization process but appeared at different trials, showing low objective values occurred at multiple stages of the optimization process. Although all barcodes were optimized within identical parameter bounds, neither the optimal objective values nor the associated parameter configurations converged to a common solution. Instead, each barcode favored a distinct region of the parameter space, resulting in barcode-specific optimization landscapes as captured by the Optuna search process and visualized in Figure R3.

#### 3.3.2 OTU Yield and Diversity Metrics (CPU)

Alpha diversity generated by the CPU-based pipeline was evaluated using the Shannon index and summarized by dominant genus using violin plots, which depict the distribution of observed values, central tendency, and samples density across diversity levels, as shown in Figure R4. Higher Shannon index values indicate increased within-sample OTU richness and evenness, whereas lower values reflect reduced diversity dominated by fewer OTUs. Differences in the vertical position and extension of the violins therefore reflect systematic differences in the magnitude and spread of alpha diversity recovered across genera.

**Figure R4.**
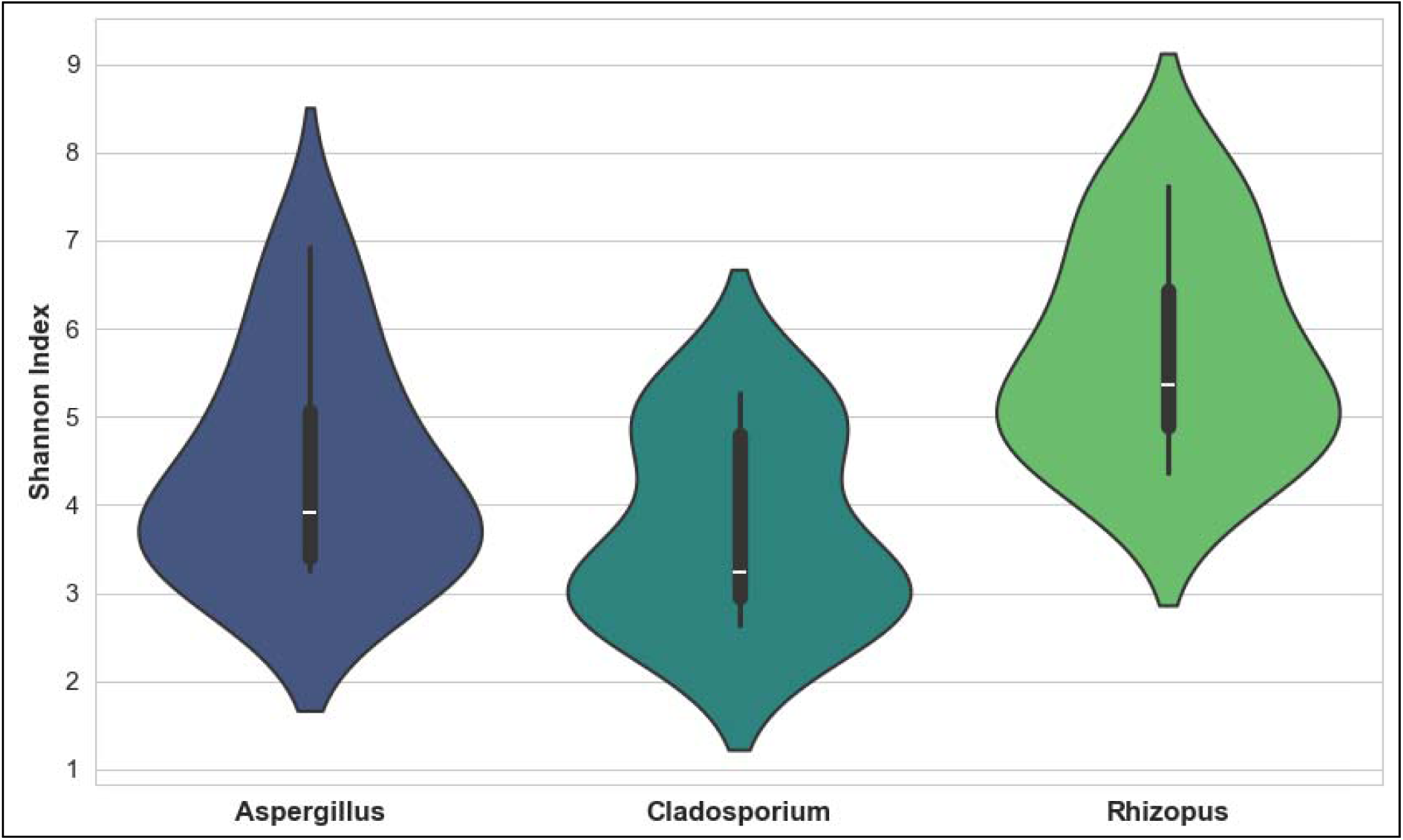
Shannon diversity per expected group

Aspergillus dominated samples exhibited intermediate Shannon diversity values, with most observations concentrated between approximately 3.3 and 4.5. The median was positioned near the center of this interval, indicating relatively consistent diversity estimates across samples. A limited number of samples exhibited Shannon values above 6 in a limited number of cases, while its greatest width occurred near the central range, reflecting a high concentration of samples with comparable diversity levels. The overall vertical extension suggests moderate dispersion in OTU diversity across samples.

Cladosporium samples displayed lower overall Shannon diversity values, spanning a range from approximately 2.5 to 5.2. The violin exhibited two distinct regions of increased width centered at those Shannon values, indicating concentration of samples around two discrete diversity levels. The median was positioned closer to the lower region, reflecting a predominance of low diversity outcomes. The comparatively shorter vertical extension relative to Rhizopus indicates a more constrained range of OTU diversity across samples.

In contrast, Rhizopus dominated samples showed substantially higher Shannon diversity values than other genera. The violin extended from approximately 4.5 to 6.5 reflecting a strong concentration of samples yielding high alpha diversity. This pattern is consistent with elevated OTU richness and evenness across Rhizopus samples.

### 3.4 GPU Pipeline Results

#### 3.4.1 Amplicon Sorter Clustering Profile

Cluster size distributions generated by the GPU-based Amplicon Sorter workflow were summarized by expected taxonomic group using violin plots, as shown in Figure R5. These plots depict the range of cluster sizes recovered, the central tendency within each group, and the concentration of clusters across size classes. The vertical extension of each violin reflects the span of cluster sizes obtained, while the width at a given value indicates the relative frequency of clusters of similar size.

**Figure R5.**
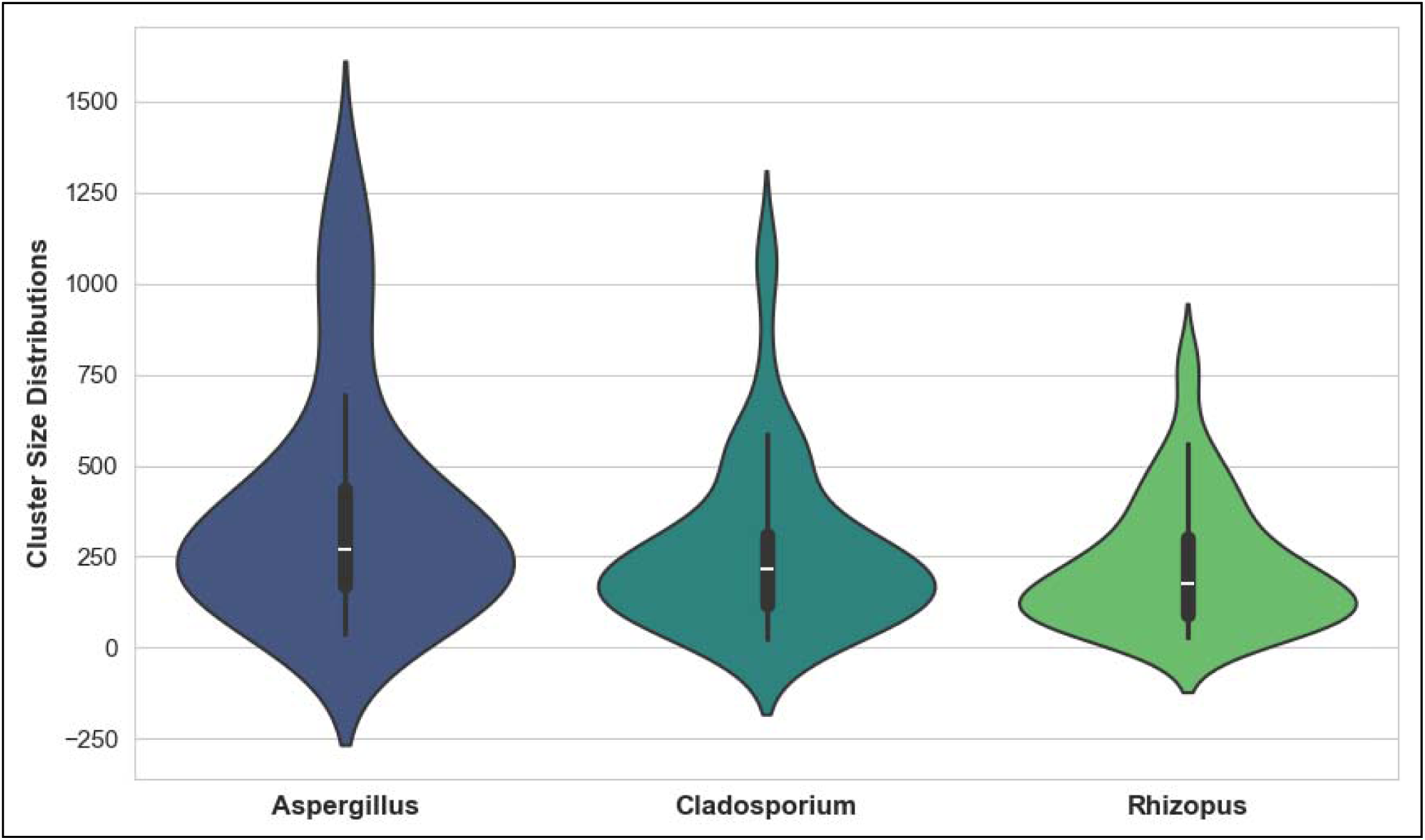
Cluster size distributions per expected group (GPU workflow)

Aspergillus-associated samples exhibited the broadest range of cluster sizes among the three groups. The distribution extended from small clusters below 100 reads to large clusters exceeding 1000 reads. The median cluster size was positioned around the mid-200 range, with the widest portion of the violin occurring between approximately 150 and 350 reads, indicating that most clusters fell within this interval. The pronounced upper tail reflects the presence of a limited number of very large clusters contributing to this distribution.

Cladosporium samples displayed a comparatively narrower size distribution. Cluster sizes were primarily concentrated between approximately 100 and 300 reads, with a median slightly above 200. The violin showed reduced extension toward larger cluster sizes compared to Aspergillus, although a smaller number of outlier clusters extended beyond 1000 reads. The greatest width of the violin occurred near the central range, indicating a strong concentration of clusters around intermediate sizes.

In contrast, Rhizopus-associated samples showed the most constrained cluster size distribution. Most clusters were concentrated at lower sizes, predominantly below 250 reads, with a median positioned near the lower end of the overall range. The vertical extension of the violin was shorter relative to other groups, and the widest region was observed between approximately 80 and 200 reads, suggesting that most clusters were small to moderately sized.

#### 3.4.2 Polishing Impact

Impact on consensus sequence quality in the GPU workflow was assessed by comparing draft consensuses generated by Amplicon Sorter with subsequent Racon and Medaka polished consensuses using pairwise alignment metrics. Across all barcodes, polishing resulted in consistent improvements in sequence identity and substantial reduction in insertion-deletion (indel) errors.

On average, draft-to-Racon comparisons yielded a mean identity of 0.966, indicating that initial consensus sequences achieved relatively high similarity to their polished counterparts. However, this stage was characterized by a high indel burden, with a mean indel rate of approximately 33.9 indels per kilobase. When Racon polished consensuses were further refined using Medaka, mean identity increased to 0.979, accompanied by a marked decrease in indel rates to an average of 20.7 indels per kilobase. This reduction represents a substantial improvement relative to both draft and Racon-only consensus.

Direct draft-to-Medaka comparisons showed a mean identity of 0.965 and an average indel rate of approximately 35 indels per kilobase, reflecting the cumulative effect of polishing while still capturing residual discrepancies inherited from draft consensus generation. Notably, identity values after Medaka polishing exhibited lower dispersion compared to earlier stages, indicating increased consistency in consensus quality across barcodes.

### 3.5 Direct Comparison of Both Pipelines

To evaluate the overall performance of the GPU and CPU pipelines in recovering expected fungal genera, taxonomic assignments were directly compared against ground-truth expectations for each barcode across the three sample groups.

**Figure R6.**
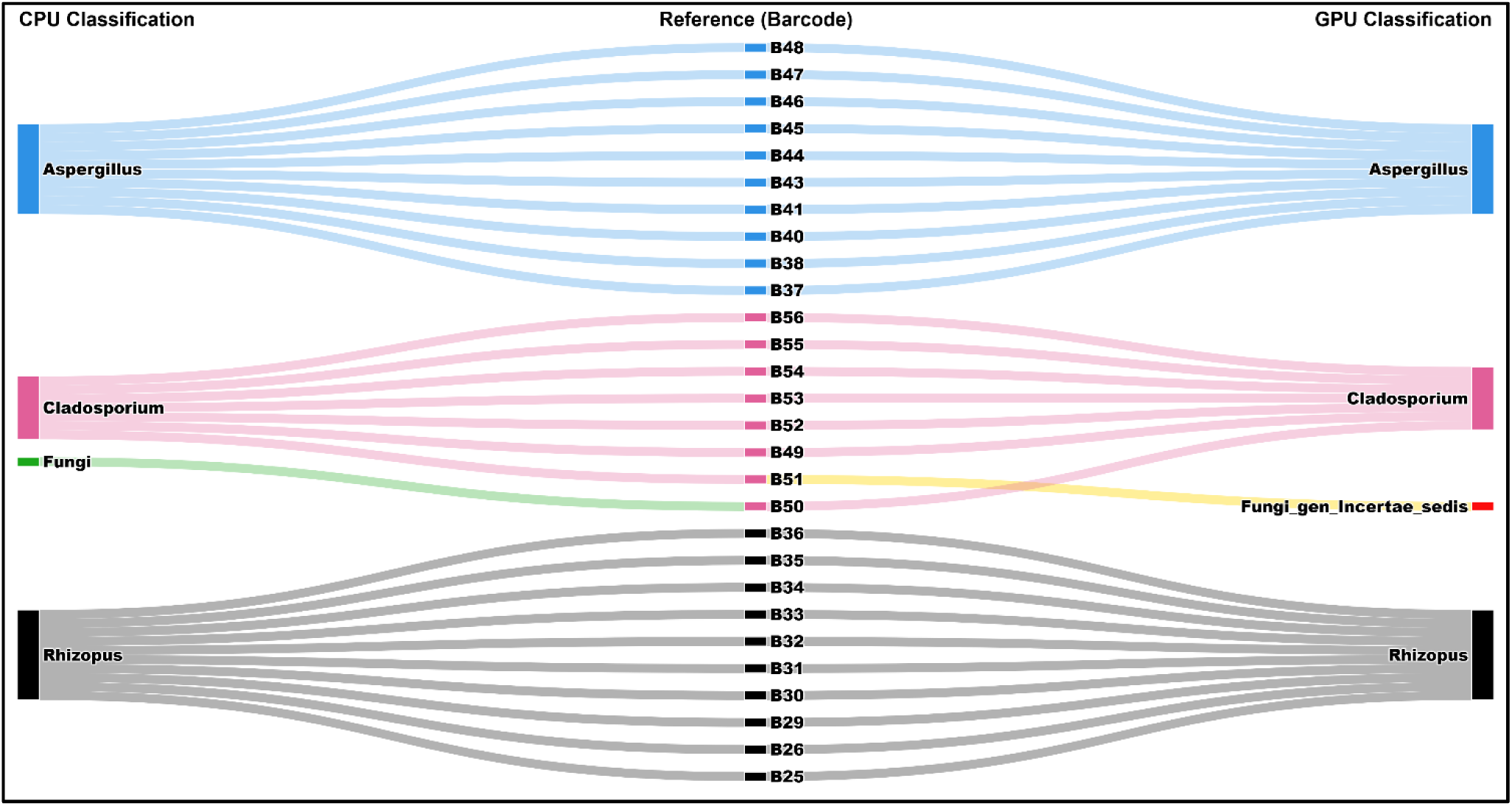
Combined Sankey Diagram (Expected taxa → CPU/GPU assigned taxa)

Figure R6 illustrates the flow from expected to assigned taxa in a combined Sankey diagram, highlighting concordance and discrepancies between pipelines. Overall, both pipelines assigned the dominant genus matching the expected group in 27 out of 28 barcodes. In the Banana group, 7 of 8 barcodes were assigned as Cladosporium by both the CPU and GPU pipelines; the remaining barcode was assigned as Fungi by the CPU pipeline (B50) and as Fungi_gen_Incertae_sedis by the GPU pipeline (B51). For the pineapple group, all barcodes were assigned as Aspergillus in both pipelines. For the Pitahaya group, all barcodes were assigned as Rhizopus in both pipelines.

Beyond genus level concordance, clear differences emerged when assignments were evaluated at finer taxonomic resolution. At the species level, the GPU polishing pipeline achieved correct assignments for 18 out of 28 barcodes, corresponding to a species-level accuracy of 64.29%. In contrast, the CPU pipeline correctly resolved species level assignments for 13 out of 28 barcodes, yielding an accuracy of 46.43%.

In terms of the total number of reported taxonomic assignments, the CPU pipeline produced a substantially larger number of total taxonomic assignments, yielding 881 genus or species level entries across all barcodes, compared to 171 assignments by the GPU polishing pipeline.

## 4 Discussion

### 4.1 Summary of Main Findings

This study compared the performance of two bioinformatics workflows for processing Nanopore sequencing ITS amplicons: a CPU-based pipeline optimized using machine learning (ML) and a GPU-accelerated pipeline with polishing based on neural networks. Using a multiplexed dataset of known expected fungal taxa, we established a benchmark to determine the taxonomic accuracy and scalability of both computational architectures.

The GPU-driven pipeline, incorporating SUP base calling models and iterative polishing algorithms, demonstrated superior ability to generate high-fidelity consensus sequences. This configuration mitigated systematic errors inherent in long-read sequencing, such as indels in homopolymeric regions, resulting in more consistent taxonomic assignments and higher species-level resolution compared to standard processing. Noise reduction in the base calling stage allowed the retention of higher-confidence data.

Conversely, the CPU-based workflow, despite using faster but less accurate basecalling models, showed a significant performance improvement thanks to the integration of machine learning-based hyperparameter optimization. The implementation of the Optuna Bayesian engine allowed for dynamic adjustment of clustering and filtering thresholds, adapting to the specific characteristics of each barcode. This approach successfully stabilized OTU recovery, validating the feasibility of robust analysis in resource-constrained infrastructures.

The performance of both pipelines varied depending on the fungal group analyzed, reflecting the influence of intrinsic biological factors, such as amplicon length and sequence complexity, on the efficiency of the processing algorithms. The GPU pipeline maximizes absolute taxonomic accuracy, while the optimized CPU pipeline offers a scalable and accessible solution, demonstrating that the choice of pipeline should be tailored to computational availability and the specific objectives of the study.

Given the complexity observed in the management of ITS regions and the need to mitigate technical noise without relying exclusively on high-performance hardware, it is therefore pertinent to delve deeper into how automated optimization influenced data quality in standard CPU-based architectures.

### 4.2 Initial reads and basecalling behavior

Table R1 shows that the GPU-based pipeline using the Super High Accuracy (SUP) model retained between 65% and 87% of the reads after quality filtering, while CPU processing with the Fast (FAST) model resulted in massive data loss, retaining only between 36% and 53%. This difference is attributed to the direct correlation between basecalling accuracy and the effectiveness of adapter demultiplexing and trimming algorithms. Fast models generate reads with lower quality scores (Q-scores) and higher error rates in terminal sequences, preventing tools like Porechop from correctly identifying adapters, resulting in the discarding of valid reads due to ambiguity or low quality (Sereika et al., 2022). While a Q10 cutoff has been adopted as a standard filtering threshold for SUP basecalling in amplicon-based microbial workflows such as 16S rRNA sequencing (Lao et al., 2023), the same threshold was deliberately applied to FAST basecalling in this study to enforce consistent quality control across models.

On the other hand, a reduction in read length was observed in the CPU pipeline compared to the GPU pipeline for the same biological samples. The median lengths in the CPU group are consistently smaller, indicative of the indel (insertion-deletion) error bias inherent in nanopore sequencing for less complex basecalling models. FAST models tend to introduce artificial deletions in homopolymer regions, shortening the final consensus sequence, while the SUP model, based on neural networks, corrects these systematic errors, recovering a sequence length closer to the actual biological size of the fragment (Delahaye & Nicolas, 2021).

On the other hand, Figure R1 reflects the technical superiority of the Super High Accuracy (SUP) basecalling model used on the GPU, which drastically reduces error rates during the conversion of the electrical signal to nucleotides. By improving base-by-base quality, the probability of sequences exceeding subsequent quality filtering thresholds is increased, avoiding the massive loss of information that characterizes the fast but imprecise models executed on the CPU (Wick et al., 2019).

The broadened morphology and lower medians observed in the violin diagrams of the CPU group demonstrate the bottleneck effect generated during the trimming and quality control stage. In rapid model-based (FAST) workflows, the high error rate in the terminal read regions hinders the algorithmic recognition of sequencing adapters, causing bioinformatics tools like Porechop to discard valid reads by misclassifying them as chimeras or low-quality sequences. Conversely, the vertical expansion and larger violin volume in the GPU group show significantly more efficient data retention; that is, the SUP model corrects indel and substitution errors, allowing for accurate adapter identification and preserving a greater proportion of the original sequenced biomass, which is critical for diversity studies where read depth directly correlates with taxonomic sensitivity (Sereika et al., 2022).

The difference observed in Figure R2 can be attributed to the neural network architecture used in each model; SUP models require massive parallel computing power due to their more complex convolutional and recurrent layers, allowing them to deconvolve the stream signal with greater fidelity and reduce stochastic noise compared to lightweight CPU models, which they cannot resolve effectively (Wick et al., 2019).

The consistent increase in quality across taxa with different genomic compositions suggests that GPU processing offers an advantage. This improvement in base-by-base accuracy is fundamental to mitigating the formation of spurious OTUs during clustering and to ensuring reliable taxonomic assignment, especially in the discrimination of closely related species where sequencing errors could be confused with real biological variability (Delahaye & Nicolas, 2021).

### 4.3 Interpretation of the Machine-Learning–Optimized CPU Workflow

The intrinsic limitations of ITS regions complicate clustering, consensus generation, and downstream taxonomic assignment, particularly when fixed or manually selected parameter thresholds are applied uniformly across samples. The selection of hyperparameter ranges was informed by the substantial methodological heterogeneity reported across fungal ITS metabarcoding studies and the need to accommodate diverse sequence characteristics inherent to this marker.

For the fungal nuclear ribosomal ITS, reported length distributions indicate that most ITS barcodes fall approximately within a 500-800 bp range (Schoch et al., 2012, p. 1). The selected conservative interval of 400-1000 bp accommodated biological variability without overly restrictive filtering. The lower bound retained complete ITS amplicons from taxa with shorter-than-average spacer regions while excluding clearly truncated reads, whereas the upper bound encompassed longer ITS variants and reduced the inclusion of anomalously long or artifactual sequences.

Minimum Phred quality thresholds vary widely in the literature dealing with Nanopore-based fungal ITS studies. Some workflows retain reads at relatively permissive thresholds, such as Q7 corresponding to default Guppy or BugSeq settings (Theologidis et al., 2023, p. 5), while others apply more stringent filtering at Q15 (Ohta et al., 2023, p. 8). This represents a balance between retaining sequencing depth while limiting error rates in inherently noisy long-read data. The explored range of Q10 - Q20 encompassed a permissive lower bound that removes extremely low-quality reads without greatly compromising coverage, and an upper bound representing common thresholds applied in post-basecalling refinement.

Clustering identity thresholds reported for fungal ITS data range from permissive to highly stringent values, depending on the targeted region and analytical objective. Ohta et al. (2023, p. 3) explicitly adapted --id thresholds by ribosomal subregion, using values as low as 0.85 for shorter ITS regions and up to 0.89 for longer ribosomal targets. In contrast, other studies applied substantially higher thresholds, such as ≥0.97 for OTU-based species identification (Langsiri et al., 2023, p. 5). Given the absence of a universally accepted ITS similarity cutoff and the documented dependence of optimal thresholds on sequencing error profiles and taxonomic scope, the broad range of 0.85-0.99 enabled systematic evaluation of clustering behavior across permissive and stringent regimes.

Minimum cluster size requirements are inconsistently reported but are often implicitly enforced through singleton filtering. Several studies explicitly discard unclustered reads or singletons, effectively imposing a minimum cluster size of two reads (Langsiri et al., 2023). Increasing the minimum number of reads required for consensus generation increases confidence in the resulting sequences but may disproportionately exclude low-abundance taxa. The explored range of 2-5 reads balanced error suppression with the retention of rare but potentially biologically relevant taxa.

Explicit positional identity thresholds for base calling from multiple sequence alignments are not standardized in fungal ITS Nanopore workflows. Most published pipelines generate consensus sequences implicitly, using centroid sequences or polishing-based approaches, without reporting numeric majority-rule cutoffs (Ohta et al., 2023). Nevertheless, general-purpose consensus tools and amplicon analysis frameworks implement explicit majority or plurality thresholds. For example, EMBOSS cons applies a default plurality cut-off of half the total sequence weight (Rice et al., 2000), while commercial software such as Geneious allows user-defined thresholds in the range of ∼0.5-0.75 for unambiguous base calling (Geneious Prime User Manual, 2025). Given the absence of standardized values and the known impact of Nanopore-specific errors such as homopolymer-associated indels, the defined consensus identity threshold range of 0.4-0.8 spanned permissive criteria that retain variable positions in noisy alignments to more stringent criteria that reduce ambiguity, enabling Optuna to empirically identify an appropriate balance for the multi-taxonomic fungal ITS dataset analyzed.

The integration of machine-learning-driven hyperparameter optimization via Optuna substantially improved the performance and robustness of the CPU based ITS processing pipeline by directly addressing well-known limitations. As demonstrated by the barcode specific optimization landscapes reported in Figure R3, Optuna stabilized clustering and polishing behavior across trials by efficiently exploring high-dimensional and non-linear parameter spaces. Parameters such as clustering identity thresholds, minimum cluster sizes, read length cutoffs, and quality filters interact in complex ways with ITS-specific sequence features. Homopolymer-associated insertion and deletion errors further amplify the sensitivity of clustering algorithms such as VSEARCH to small changes in thresholds values (Calus et al., 2018).

By iteratively refining parameter choices based on the observed objective values across trials, the optimization process converged on configurations that minimized infeasible outcomes while adapting to barcode specific sequence characteristics, as reflected by the differing proportions of infinite and finite objective values across barcodes in Figure R3.

The observed barcode dependent optimization landscapes directly reflect this stabilization effect. Barcodes differed not only in the magnitude of their optimal objective values but also in the trial at which these optima were reached, as shown by the trial-value trajectories in Figure R3, indicating that Bayesian optimization did not simply identify a universal parameter set but instead adapted to the underlying heterogeneity of the ITS data. This behavior is consistent with the expectation that ITS variability, including differences in read length distribution and motif composition, requires flexible rather than fixed parameterization to achieve reliable clustering outcomes.

A key advantage of the Optuna-based approach is its ability to reduce subjective human bias, which remains a pervasive issue in ITS workflows. Manual parameter tuning is often guided by heuristics, prior experience, or dataset-specific trial and error, resulting in pipelines that are difficult to reproduce and compare across studies (Edgar, 2010). The distinct optimal parameter combinations identified for each barcode in Figure R3 illustrate how automated optimization replaces subjective decision making with data driven parameter selection derived directly from dataset behavior. This demonstrates that automated tuning can capture sample specific requirements that would be difficult to anticipate manually.

Group specific patterns further highlight the biological relevance of adaptative optimization. Samples dominated by Rhizopus exhibited higher sensitivity to clustering thresholds, consistent with known intragenomic ITS variation and highly variable ITS2 regions in this genus. Such characteristics are known to increase susceptibility to chimera formation and misclustering when stringent identity thresholds are applied (Woo et al., 2010). This heightened sensitivity is congruent with the broader range of optimal objective values and parameter configurations observed for Rhizopus associated barcodes in Figure R3, as well as with the elevated and widely distributed Shannon diversity values reported in Figure R4. In contrast, Aspergillus dominated samples displayed more stable clustering behavior, reflected both in narrower objective value distributions in the optimization landscape in Figure R3 and in more tightly centered Shannon diversity values in Figure R4. These differences illustrate how machine learning driven optimization can implicitly accommodate taxon specific sequence properties without requiring explicit taxonomic information during parameter selection.

Also, the use of automatic hyperparameter selection confers important reproducibility advantages. By fixing the random seed, logging optimization trajectories, and storing complete study states, Optuna enables exact replication of optimization outcomes and transparent reporting of parameter choices. The explicit reporting of barcode specific optimization trajectories and diversity outcomes in Figures R3 and R4 exemplifies this reproducibility, linking parameter selection directly to observable clustering and diversity metrics. This addresses a longstanding limitation in bioinformatics workflows, where undocumented manual adjustments often impede reproducibility and cumulative analysis. In applied contexts such as environmental monitoring, agricultural surveillance, and post-harvest fungal diagnostics, these advantages support the development of scalable and standardized ITS workflows that remain sensitive to biological heterogeneity while maintaining consistency and interpretability.

### 4.4 Interpretation of the GPU Polishing Workflow (Medaka + Racon)

The improvements observed in consensus sequence accuracy within the GPU workflow can be attributed to the complementary roles of Racon and Medaka in correcting systematic error patterns characteristic of nanopore-based ITS sequencing. While Amplicon Sorter provides an efficient clustering and draft consensus generation step, as reflected in the cluster size distributions shown in Figure R5, residual base-level inaccuracies remain and are subsequently addressed through multistage polishing.

Racon primarily reduces systematic alignment-level inconsistencies by leveraging read to consensus alignments, resulting in substantial reductions in indel burden relative to draft consensuses. Medaka further refines these sequences using a neural network trained on nanopore signal characteristics, enabling the correction of systematic errors that persist after alignment-based polishing. The quantitative improvements observed after polishing, including sequence identity and marked reductions in indel rates across barcodes, indicate that polishing contributes not only incremental corrections but also enhanced consistency in final consensus sequences, as reflected by the reduced dispersion of identity values following Medaka refinement.

The effectiveness of polishing varied across fungal groups, reflecting an interaction between biological ITS variability and error-correction dynamics. Although Aspergillus-associated samples typically formed larger clusters with higher read coverage as shown in Figure R5, improvements in consensus accuracy did not consistently translate into enhanced species-level resolution, likely reflecting the limited discriminatory power of ITS regions among closely related Aspergillus species. In contrast, Rhizopus-associated samples, despite exhibiting more constrained cluster size distributions and higher intragenomic ITS variability, showed a more pronounced improvement in species-level assignments following polishing. This suggests that, for taxa characterized by heterogenous ITS regions, like Rhizopus (Woo et al., 2010), GPU-based polishing primarily acts to suppress nanopore-specific technical noise rather than collapsing genuine biological variation, thereby enabling more accurate species discrimination. These observations highlight that polishing effectiveness is not solely determined by read depth or cluster size, but by the extent to which error correction enhances biologically informative signal relative to intrinsic sequence variability across taxa.

### 4.5 Comparative Interpretation of CPU–ML vs GPU–Polishing

The Sankey diagram illustrates a high degree of concordance at the genus level between the CPU and GPU pipelines, with agreement observed in 27 out of 28 barcodes. This alignment reflects the robustness of both approaches in capturing the dominant fungal genera expected from the samples, with discrepancies limited to one instance. Such conservative fallbacks likely reflect cases where sequence variability or limited taxonomic signal challenges precise genus delineation, yet the overall genus level consensus underscores that both pipelines effectively resolve the core taxonomic structure of the fungal communities under study.

Despite this genus level harmony, the pipelines diverge notably at the species level, as evidenced by the quantitative metrics showing a slightly superior species level accuracy in the GPU pipeline compared to the CPU pipeline. This species level divergence coexists with genus agreement because the pipelines differ in their handling of intra-barcode sequence diversity, leading to distinct resolutions of species boundaries. The majority species analysis per barcode further elucidates this pattern: GPU outputs typically consolidate taxonomic assignments into a single dominant species per barcode, fostering confident species assignments by minimizing ambiguity from variant sequences. In contrast, CPU outputs frequently retain multiple competing species level assignments within the same barcode, which dilutes dominance and results in broader but less precise taxonomic profiles.

These differences manifest in the pronounced contrast in assignment abundance, with the CPU pipeline yielding 881 total assignments across barcodes versus 171 for the GPU pipeline. The higher abundance in CPU outputs suggests a tendency to preserve a wider array of potential taxa, which reflects a tendency to preserve a wider range of potential taxa, accommodating subtle sequence variations but at the cost of species-level precision. Conversely, the GPU’s lower abundance indicates a more stringent consolidation process, prioritizing a dominant species signal that aligns more closely with expected taxa in most barcodes. The GPU pipeline homogenizes taxonomic profiles through better consensus formation, while the CPU pipeline preserves within group diversity signatures, potentially capturing low abundance variants that contribute to ecological insights.

Observed misclassifications can be attributed to multiple sources, including PCR bias in ITS amplicon sequencing, which may preferentially amplify certain variants and distort community representation (Bellemain et al., 2010), intragenomic ITS variation introducing heterogeneous copies within genomes that complicate species delineation (Lindner & Banik, 2011), and limitations of reference databases, such as incomplete fungal ITS coverage leading to unassigned or erroneous taxa.

To minimize such errors, the hybrid taxonomic assignment approach combined the high-local alignment capabilities of BLAST for ITS regions with the probabilistic syntactic classification framework of SINTAX, enabling more robust genus- and species-level assignments compared to either method alone. The threshold values applied in the decision rules were informed by empirically validated performance benchmarks for fungal ITS identification.

For species-level assignments, Edgar (2018a, p. 18) assessed SINTAX performance on fungal ITS datasets and demonstrated that species-level assignment rapidly loses precision below bootstrap values of 0.8, recommending higher thresholds for conservative classification. Accordingly, a SINTAX species confidence threshold of 0.9 was adopted for high-confidence species identification. In parallel, BLAST-based species assignment was constrained by a minimum percent identity of 97%, a threshold commonly applied for fungal ITS species-level identification (Raja et al., 2017). Raja et al. (2017, p. 6) further recommend minimum query coverage values of approximately 80% for reliable ITS-based species identification, noting that partial alignments restricted to conserved subregions can inflate percent identity and lead to spurious assignments. In the present study, this coverage threshold was increased to 90% to enforce near full-length ITS alignment, thereby reducing the influence of short high-identity matches and improving robustness in Nanopore-derived reads, which are more prone to localized errors and length heterogeneity.

Cases of extreme BLAST dominance were defined to capture instances where a single reference sequence overwhelmingly outperformed all alternatives. Vu et al. (2019, p. 9) demonstrated through in-depth analysis of validated fungal ITS sequences that species discrimination by ITS requires almost perfect sequence similarity, with an estimated optimal species boundary of approximately 99.6% sequence identity. A conservative BLAST identity threshold of ≥99.0% was therefore established to identify near-unambiguous species-level matches, even when probabilistic classifier support was unavailable or inconclusive. To ensure that high identity values reflected biologically meaningful alignments rather than short local matches, a high alignment coverage requirement of ≥95% was imposed, consistent with the full-length sequence comparisons underlying the similarity analyses reported by Vu et al. (2019, p. 14). Additionally, a minimum percent identity difference of ≥1.0% between the top and second BLAST hits was required to confirm that the top hit was decisively superior, reflecting clear separation rather than a cluster of closely related taxa with comparable similarity scores.

Genus-level assignments were supported by concordant probabilistic and alignment-based evidence. Vu et al. (2019, p. 12) demonstrated, based on a large, curated dataset of filamentous fungal reference strains, that genus-level discrimination using ITS sequences is optimally achieved at sequence similarity values around 94.3%. This empirically derived boundary was adopted as a reference point, and a BLAST percent identity threshold of ≥94.0% was applied as an approximate and conservative operational threshold for genus-level assignments. Winand et al. (2025) further showed that a minimum BLAST query coverage of approximately 80% is required to retain reliable taxonomic assignments at the genus level in fungal ITS metabarcoding, as lower coverage values increase the likelihood of partial or spurious matches. In parallel, Edgar (2018b, p. 18) evaluated SINTAX classifier performance on fungal ITS datasets and showed that genus-level assignments retain acceptable accuracy at bootstrap confidence values around 0.8, while higher thresholds substantially reduce the risk of over-classification. Consequently, a stricter SINTAX genus confidence threshold of ≥0.90 was applied to prioritize precision over recall in genus-level consensus calls.

Group specific patterns reveal why certain expected taxa were more readily recovered by one pipeline over the other. For Rhizopus stolonifer, the CPU and GPU pipelines achieved a perfect recovery (10/10 barcodes), suggesting that, despite high intragenomic ITS variability, the dominant species signal in Rhizopus stolonifera is sufficiently strong to be recovered consistently by both pipelines (Woo et al., 2010). In contrast, for Aspergillus niger, the GPU pipeline excelled (7/10 correct versus 3/10 in CPU), consistent with the ability of consensus-based polishing to mitigate intragenomic ITS variation known in this group, reducing misclassifications to closely related species like Aspergillus tubingensis or ambiguous Aspergillus sp. (Steenwyk et al., 2024). Similarly, for Cladosporium cladosporioides, the GPU recovered more expected taxa (2/8 versus 0/8 in CPU), as this complex harbor high cryptic diversity and intragenomic differences in ITS copies, which the GPU’s homogenization resolves better, avoiding frequent shifts to pseudocladosporiorides, tenuissimim, or unspecified sp. (Bensch et al., 2010).

At the barcode level, these emergent taxonomic decisions imply that the CPU pipeline is more flexible, accessible, and tunable, making it better suited for high diversity or noisy groups where preserving variants is advantageous. The GPU polishing pipeline offers higher species-level accuracy but requires more resources and relies on high read depth to achieve superior consensus.

Beyond methodological performance, the contrasting behaviors observed between pipelines carry direct biological and applied implications for fungal ITS metabarcoding. The ability of the CPU workflow to preserve within group diversity while maintaining robust genus level assignments suggests practical suitability for exploratory or surveillance-oriented applications where capturing intragenomic and population level variability is often more informative than enforcing strict consensus. Conversely, the higher species level correctness achieved by the GPU polishing pipeline highlights its relevance for high precision contexts, where minimizing sequencing induced artifacts and resolving closely related taxa are critical. This comparative analysis highlights that pipeline selection should still be guided by sequencing depth, computational constraints, and analytical objectives to maximize interpretability and reliability in applied and research contexts.

### 4.6 Comparison with Previous Work

Previous investigations into nanopore-based ITS metabarcoding for fungal identification have largely focused on individual components of the analytical workflow rather than on fully integrated, end-to-end processing strategies. Dierickx et al. (2024b) implemented VSEARCH based clustering and SINTAX taxonomic assignment as part of a fungal metabarcoding workflow, demonstrating the applicability of these tools for long read ITS analysis but without evaluating their interaction with consensus polishing steps or alternative pipeline configurations. In parallel, (Huang et al., 2021) developed Homopolish as a method to correct systematic sequencing errors in nanopore reads through homologous polishing, focusing on error-correction performance rather than on amplicon clustering or taxonomic benchmarking.

Several studies have highlighted the limitations of such component focused approaches. Loit et al. (2019) analyzed technical biases inherent to ITS metabarcoding, including chimera formation and preferential amplification, and emphasized how these factors can distort inferred fungal composition when analytical steps are considered in isolation. Similarly, Yu et al. (2023) presented a targeted nanopore sequencing pipeline applicable to organism identification yet did not extend their analysis to comparative benchmarking of preprocessing, clustering, and consensus formation within a unified end to end framework. These studies illustrate a fragmented landscape in which individual tools are validated, but their combined effects on taxonomic accuracy remain insufficiently explored in integrated workflows.

Basecalling accuracy has been addressed in a separate body of literature. Delahaye & Nicolas (2021) explicitly described the tradeoff between FAST and high accuracy basecalling modes in nanopore sequencing, linking improved accuracy to increased computational cost. Building on this, Ahsan et al. (2024) discussed practical constraints associated with GPU based processing, noting that while GPUs enable faster and more accurate basecalling, their cost and limited availability pose barriers for many laboratories, reinforcing the need for alternatives that balance accuracy and accessibility.

To date, we are not aware of published studies that have applied machine learning-driven hyperparameter optimization, such as Bayesian approaches implemented through frameworks like Optuna, specifically to ITS clustering in nanopore based fungal metabarcoding. Likewise, direct benchmarking of CPU-optimized and GPU accelerated ITS workflows using multi-genus fungal datasets with known expected taxonomic composition has not been reported. The present study addresses these gaps by integrating ML optimized clustering on CPU resources with GPU based polishing into a single, systematically evaluated framework, thereby providing a novel benchmark for nanopore ITS analysis in fungal taxonomy.

### 4.7 Strengths and Limitations

A major strength of this study is the end-to-end benchmarking of complete CPU- and GPU-based nanopore ITS workflows using a defined multi-genus dataset spanning 28 independent barcodes with known expected taxa. This design enables robust performance assessment across replicate sequencing units, capturing variability arising from read depth, consensus formation, and computational architecture, rather than from isolated algorithmic components evaluated in isolation. The integration of machine learning-guided hyperparameter optimization further strengthens the CPU-based pipeline by reducing heuristic parameter selection and improving clustering robustness across barcodes.

Another strength lies in the biologically informed interpretation of results, explicitly accounting for known properties of fungal ITS regions, including intragenomic variation and taxonomic complexity, which strengthens the biological plausibility of the comparative conclusions.

Several limitations should also be acknowledged. The benchmarking was performed on defined fungal compositions rather than standardized mock communities or highly complex environmental samples, which may exhibit additional sources of bias related to uneven abundance distributions, chimera formation, and biological complexity. Performance was evaluated using a specific set of basecalling, polishing, and clustering tools, and alternative software or future updates could influence relative outcomes. In addition, computational performance remains dependent on hardware availability, particularly for GPU based workflows, which may limit applicability in resource-constrained laboratory settings.

### 4.8 Future Directions

Future work will focus on extending the proposed framework toward greater scalability, generalizability, and methodological flexibility across diverse fungal metabarcoding scenarios. A natural next step is the development of explicitly hybrid pipelines that dynamically allocate computational tasks between CPU and GPU resources, building on the complementary strengths demonstrated in this study. In such architectures, CPU-based workflows would continue to support clustering, parameter optimization, and decision-making stages, while GPU acceleration would be selectively applied to error correction and polishing steps where performance gains are most pronounced. These adaptive strategies would facilitate efficient deployment across heterogeneous computing environments while maintaining robust and reproducible taxonomic inference.

Beyond hyperparameter optimization, the integration of additional machine learning approaches represents a promising avenue for further methodological refinement. Supervised or semi-supervised models could be explored to inform clustering thresholds, confidence scoring, or consensus selection using read-level features, barcode complexity metrics, or internal consistency signals derived directly from the data. Such approaches could enhance robustness in low-depth or high-diversity datasets, where fixed heuristics or manually selected parameter may be suboptimal, while preserving the transparency and reproducibility required for comparative metabarcoding studies.

### 4.9 Conclusion

This study establishes a systematic benchmark for nanopore-based ITS amplicon analysis, demonstrating that the integration of machine learning-driven optimization substantially enhances CPU-based workflows and narrows the performance gap with GPU-accelerated approaches. Although GPU workflows maximize consensus accuracy and species-level resolution, the dual-pipeline comparison provides practical and actionable guidance for selecting analysis strategies according to accuracy requirements and available computational resources. Overall, this work contributes a reproducible, scalable, and biologically coherent framework for nanopore ITS metabarcoding, supporting accessible analytical solutions without compromising taxonomic reliability.

## 6 Author Contributions

DA conceived and designed the study, led project administration and team coordination, developed both bioinformatics pipelines from raw FASTQ processing through final consensus generation, implemented the machine learning–driven hyperparameter optimization framework using Optuna, developed the hybrid BLAST+SINTAX taxonomic assignment module, and contributed to writing sections of the manuscript. PM performed raw signal processing and basecalling from FAST5 to FASTQ format using Dorado, contributed to writing sections of the manuscript, and organized the supplementary materials. PZ generated all data visualizations, performed quality analysis across pipelines, and contributed to writing sections of the manuscript. JR provided methodological guidance and contributed critical review and editorial suggestions throughout the manuscript development. EV supervised the research, reviewed the methods and results for scientific rigor, and provided overall direction as corresponding author. All authors read and approved the final manuscript.

## Supporting information

Figure R3 2 B34 Vsearch vs Cluster Size

Figure R3 1 B34 Vsearch vs Identity

Figure R3 4 B41 Vsearch vs Cluster Size

Figure R3 3 B41 Vsearch vs Identity

Figure R3 6 B56 Vsearch vs Cluster Size

Figure R3 5 B56 Vsearch vs Identity

## Acknowledgments

The authors gratefully acknowledge the Department of Food Science and Biotechnology (DECAB) at the Escuela Politécnica Nacional for providing access to the high-performance computing resources and databases used in this research. We express our sincere gratitude to PhD Edwin Rafael Vera Calle for his constant support, promotion of this project, and valuable time dedicated to reviewing the manuscript during its development. We also thank Mgtr. Joselyn Riczury Olmos Tovar for her expert collaboration and key insights regarding data analysis.

