## Supplementary figures and images for "Machine Learning–Enhanced Nanopore ITS Analysis: Evaluating CPU–GPU Pipelines for High-Accuracy Fungal Taxonomic Resolution"

### Figure R3 1 B34 Vsearch vs Identity

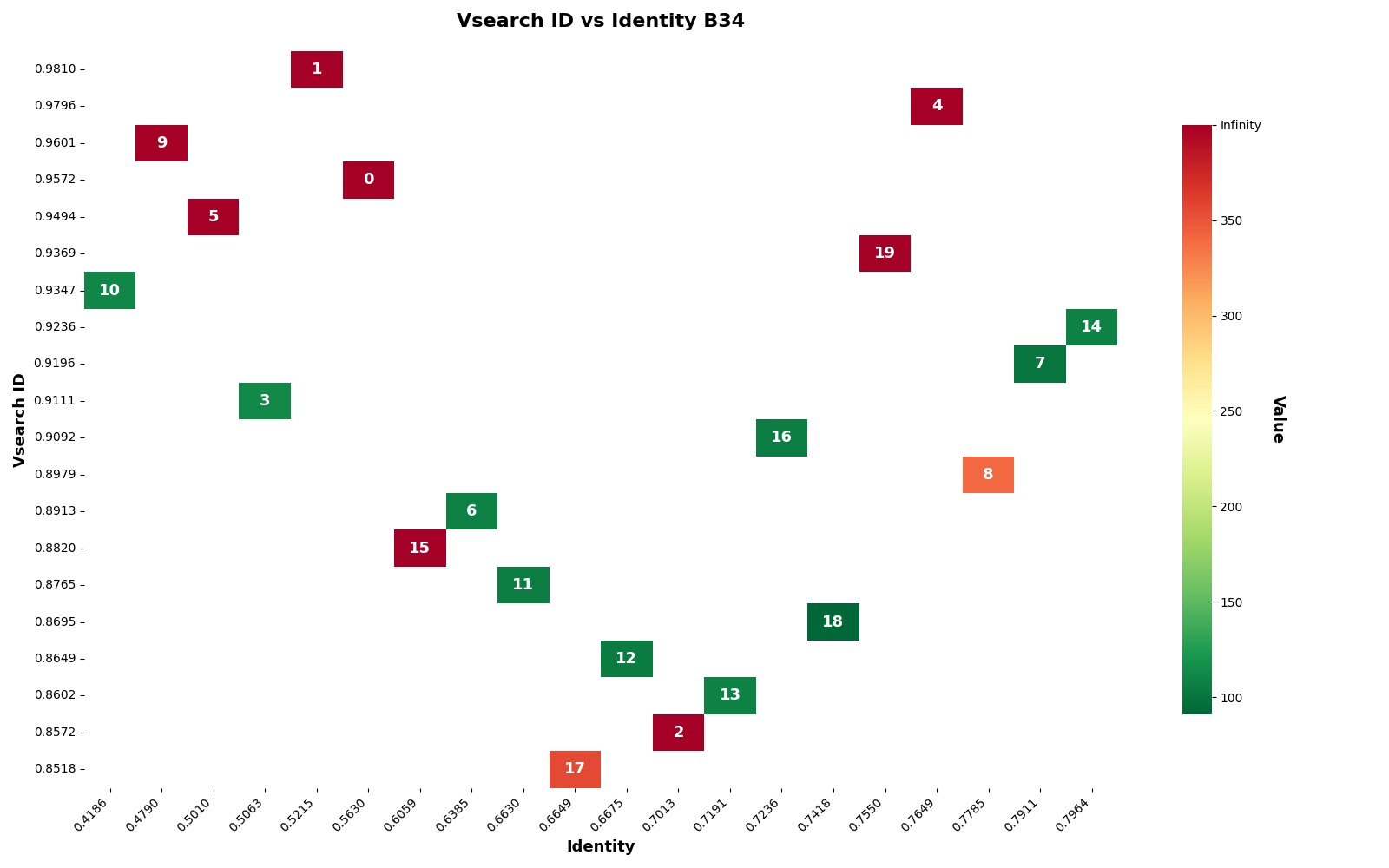

### Figure R3 2 B34 Vsearch vs Cluster Size

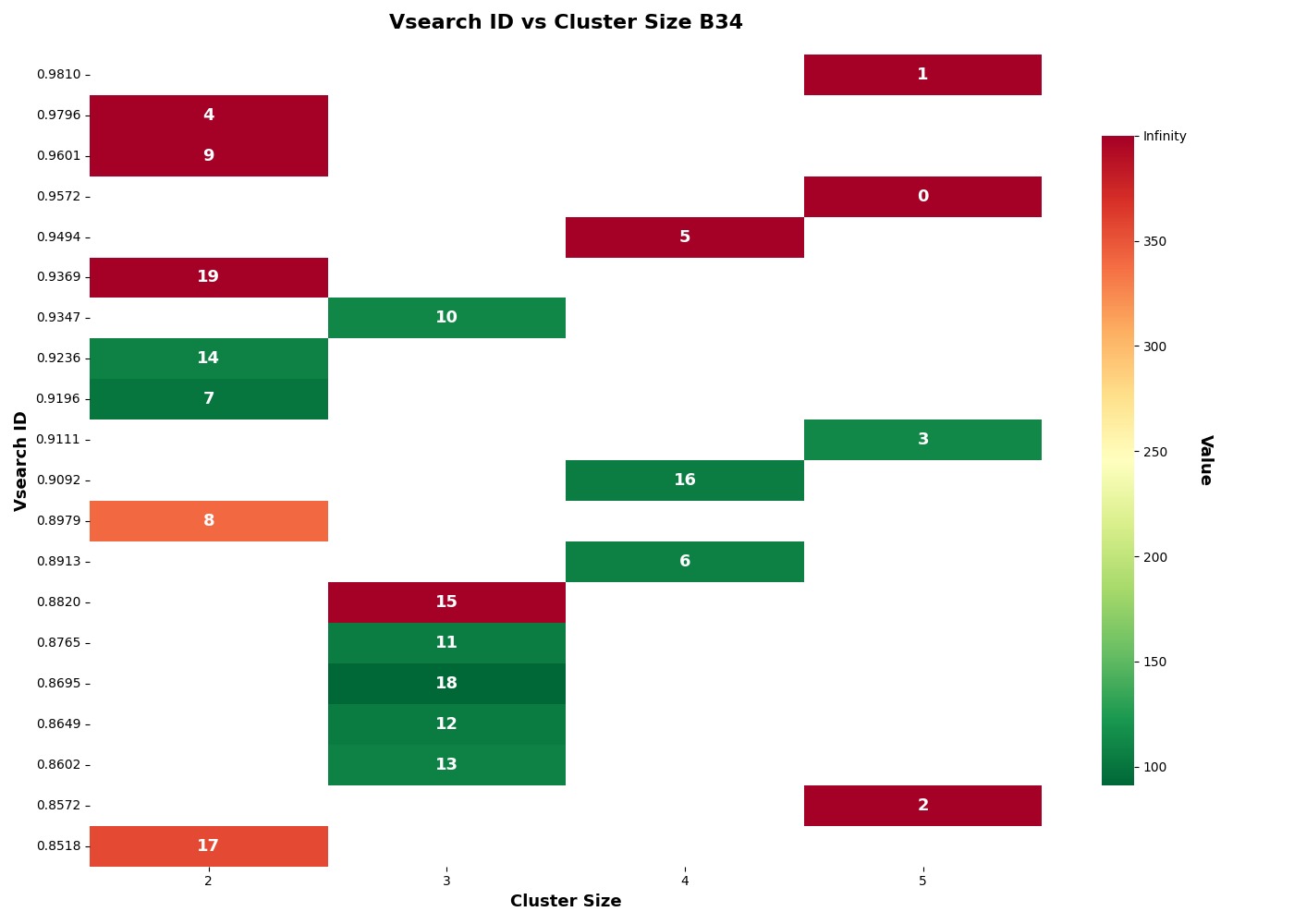

### Figure R3 3 B41 Vsearch vs Identity

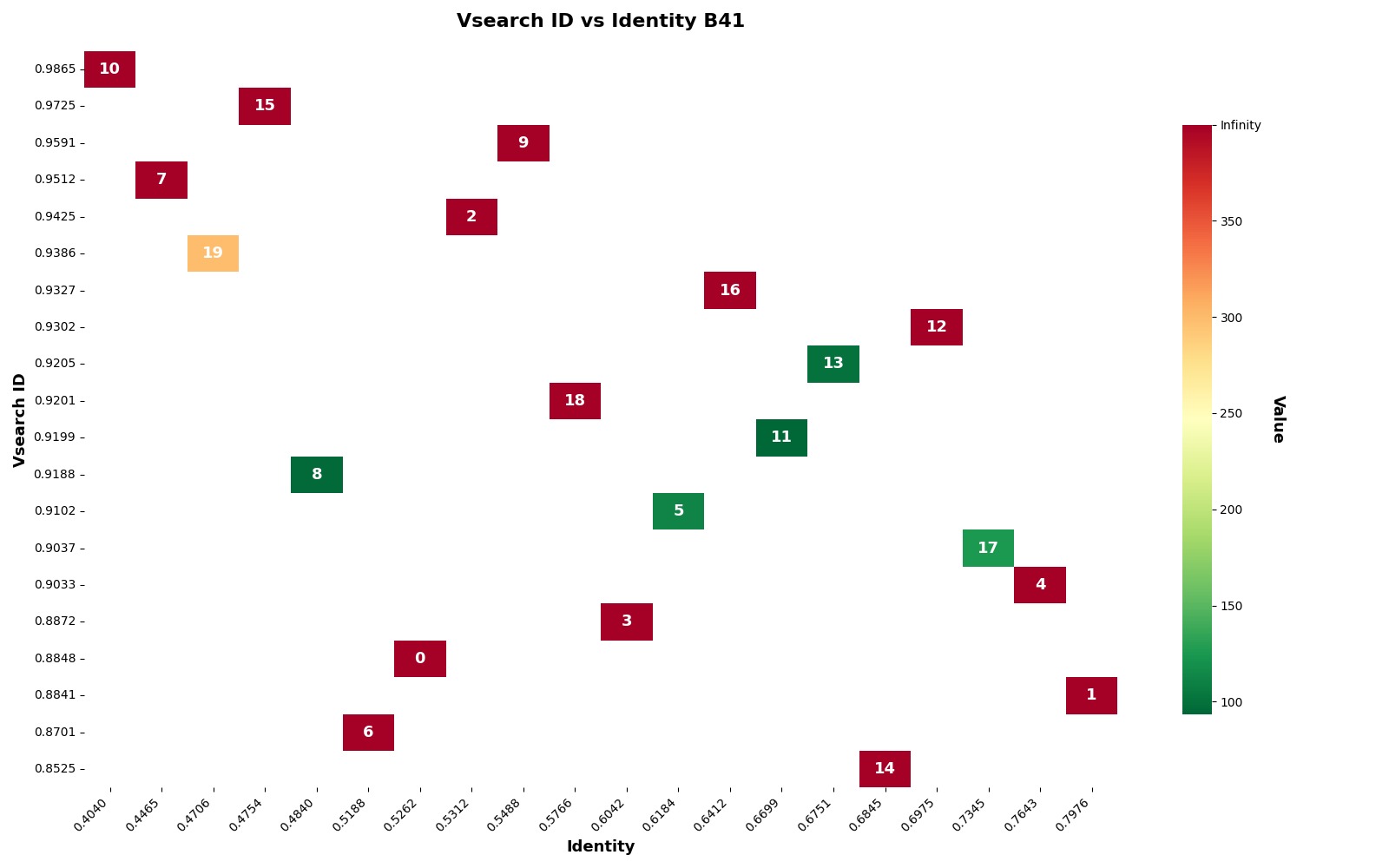

### Figure R3 4 B41 Vsearch vs Cluster Size

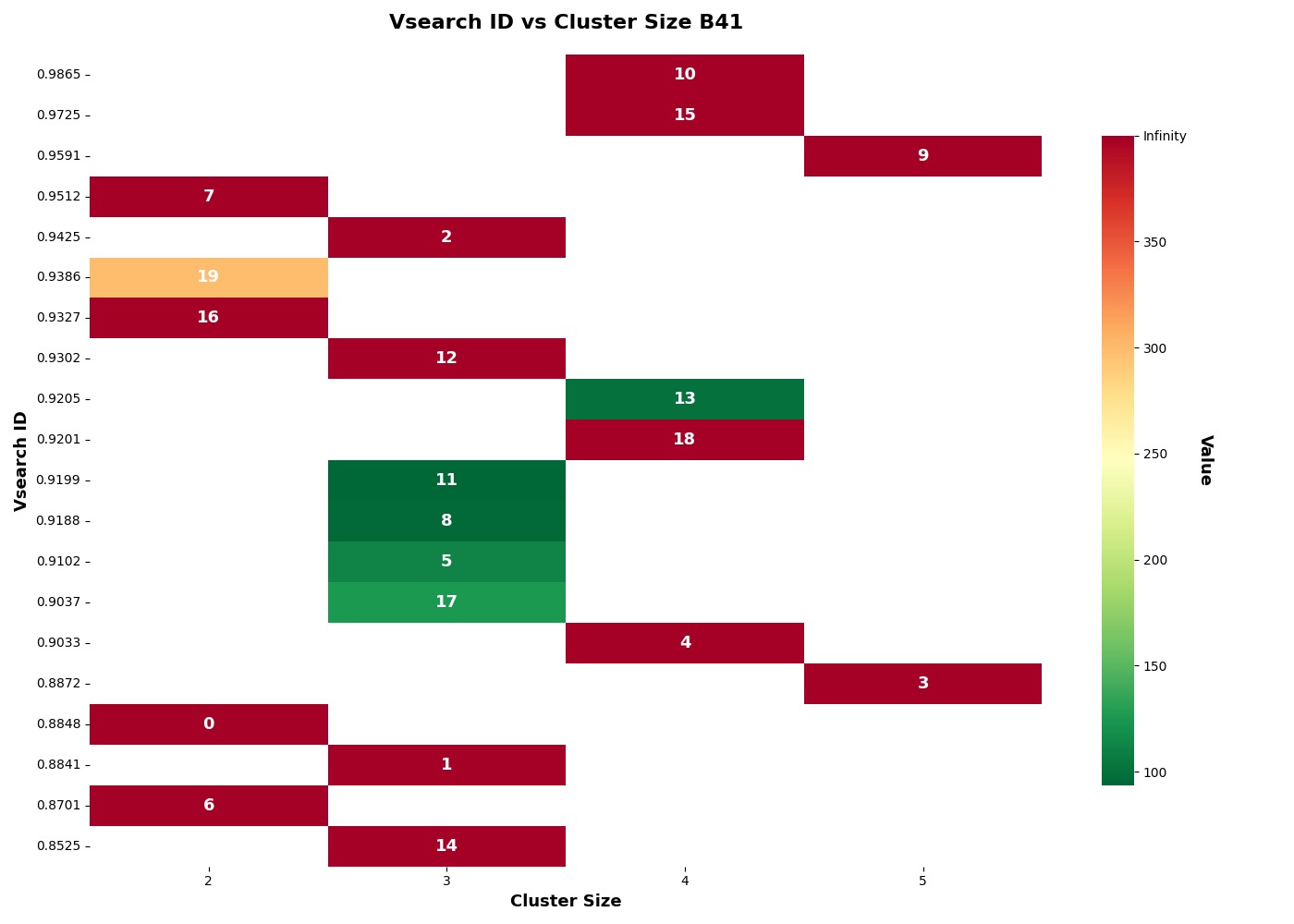

### Figure R3 5 B56 Vsearch vs Identity

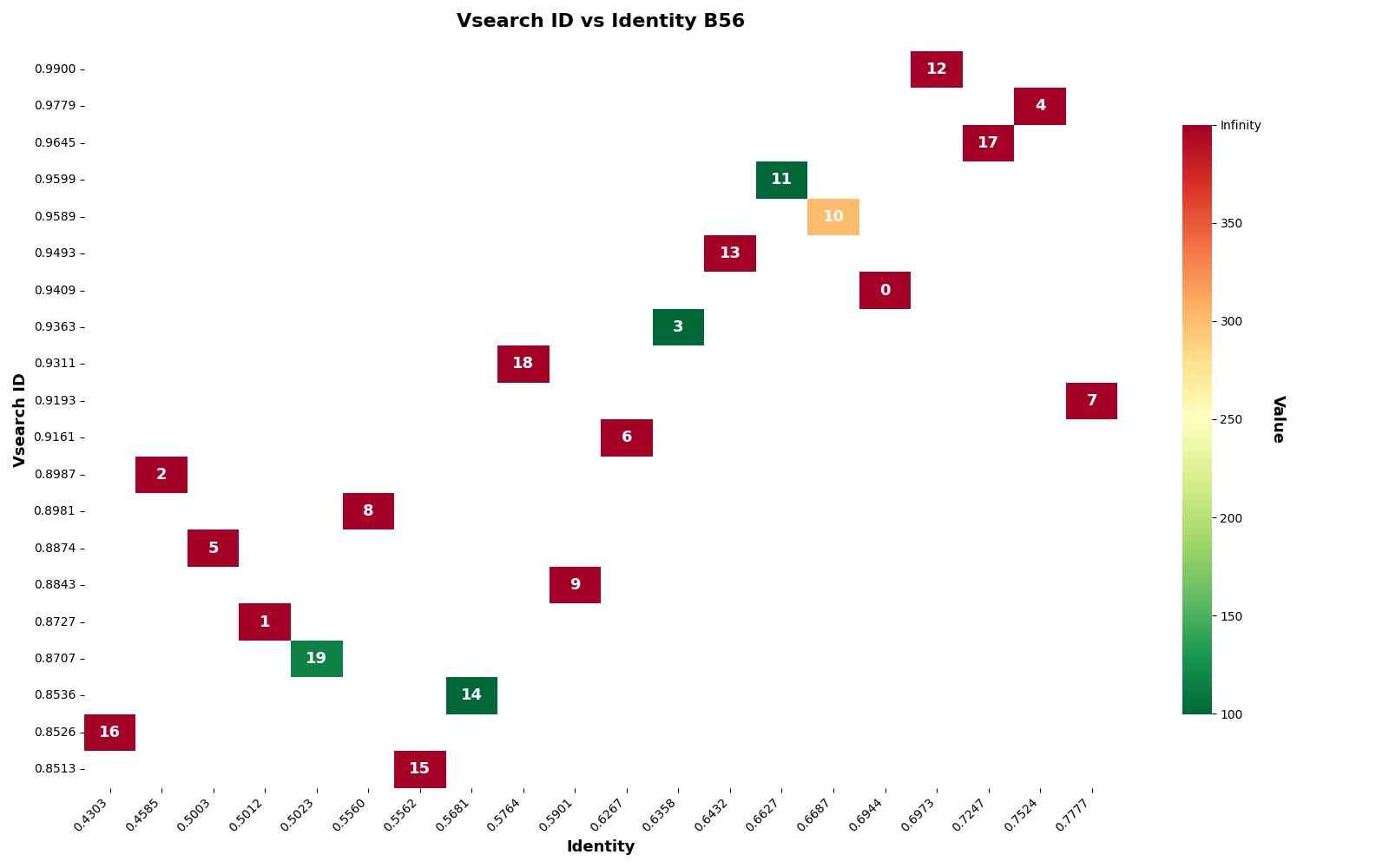

### Figure R3 6 B56 Vsearch vs Cluster Size

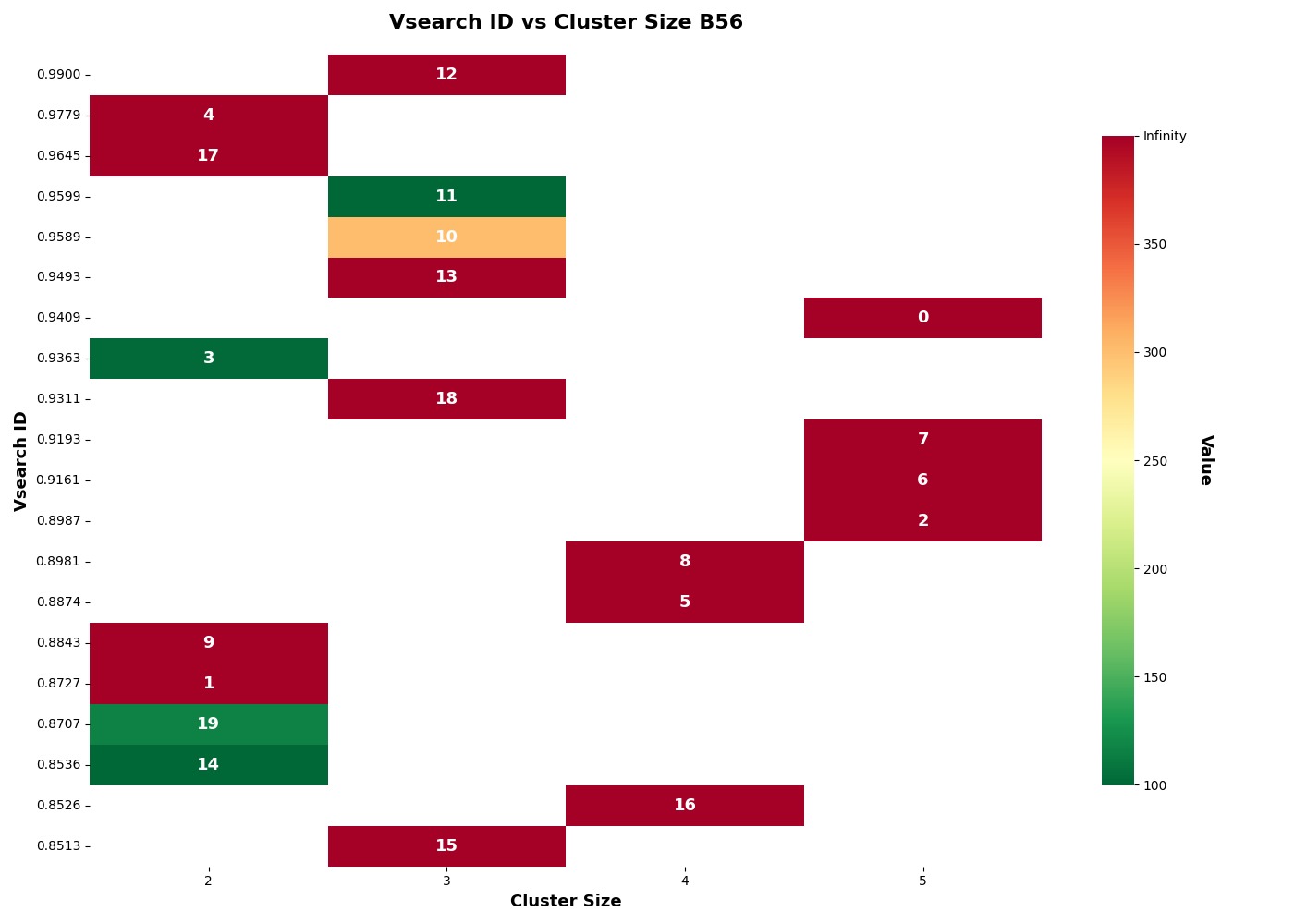
